# Urn models for regulated gene expression yield physically intuitive solutions for probability distributions of single-cell counts

**DOI:** 10.1101/2020.02.09.940452

**Authors:** Krishna Choudhary, Atul Narang

## Abstract

Fitting the probability mass functions from analytical solutions of stochastic models of gene expression to the count distributions of mRNA and protein molecules in single cells can yield valuable insights into mechanisms of gene regulation. Solutions of chemical master equations are available for various kinetic schemes but, even for the models of regulation with a basic ON-OFF switch, they take complex forms with generating functions given as hypergeometric functions. Gene expression studies that have used these to fit the data have interpreted the parameters as burst size and frequency. However, this is consistent with the hypergeometric functions only if a gene stays active for short time intervals separated by relatively long intervals of inactivity. Physical insights into the probability mass functions are essential to ensure proper interpretations but are lacking for models of gene regulation. We fill this gap by developing urn models for regulated gene expression, which are of immense value to interpret probability distributions. Our model consists of a master urn, which represents the cytosol. We sample RNA polymerases and ribosomes from it and assign them to recipient urns of two or more colors, which represent time intervals with a homogeneous propensity for gene expression. Colors of the recipient urns represent sub-systems of the promoter states, and the assignments to urns of a specific color represent gene expression. We use elementary principles of discrete probability theory to derive the solutions for a range of kinetic models, including the Peccoud-Ycart model, the Shahrezaei-Swain model, and models with an arbitrary number of promoter states. For activated genes, we show that transcriptional lapses, which are events of gene inactivation for short time intervals separated by long active intervals, quantify the transcriptional dynamics better than bursts. Our approach reveals the physics underlying the solutions, which has important implications for single-cell data analysis.

## Introduction

Gene expression occurs in multiple steps [1]. The biochemical mechanisms of its steps are of great interest [2–4]. In particular, a majority of studies have focused on gene regulation, transcription, and translation [5]. Genes might be expressed at a uniform rate or transition between two or more states with different rates of expression [6]. In the latter case, the transition kinetics might be regulated by gene-specific mechanisms such as interactions of the promoters with specific transcription factors or gene-independent mechanisms such as DNA supercoiling [7–13]. When genes are in transcriptionally active states, mRNA molecules might be produced, which might be translated further into proteins. Experimental data for the distribution of mRNA/protein molecules in single cells could be harnessed for model selection out of a candidate set of mechanistic models [14–16]. In this direction, numerous stochastic models of gene expression have been developed to study a range of kinetic schemes [5, 17–21]. Their analytical solutions for the probability distributions of molecular counts have been obtained in many cases [16,22–43], and comparisons with single-cell RNA-seq and single-molecule imaging data have facilitated inferences in mechanistic studies [6, 11, 23, 44–50].

An elementary model of constitutive gene expression uniformly allows transcription at all times [36, 37]. This is identical to the classical *birth-and-death* process, which results in the Poisson distribution for mRNA molecules at stationary state [51]. A physically intuitive method to derive the mRNA distribution utilizes an *urn model*, whereby the kinetic scheme of mRNA production (which, say, occurs with rate constant *v*_0_) and degradation (say, with rate constant *d*_0_) is mapped to an urn scheme. To this end, one considers an urn with balls of two colors —black and white. As time progresses, we sample balls one-at-a-time from the urn, i.e., we perform Bernoulli trials [51]. The outcome of each trial, a black or white ball, corresponds to an outcome of transcription or no transcription in physical terms, respectively. Each trial consists of drawing a ball, recording its color, replacing the ball in the urn, and mixing the urn to prepare for the next trial. Let us say that we draw balls without taking any break and the time duration per trial, Δ*t* is infinitesimal. The kinetic scheme is mapped to the urn scheme by defining that the proportion of black balls in the urn is the same as the probability of transcription in Δ*t* time, which is *v*_0_Δ*t*. Finally, the probability of observing *m* copies of mRNA molecules is obtained as the probability of drawing *m* black balls in infinitely many trials during the mean lifetime of mRNAs, 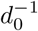. This is done by appealing to intuitive principles of combinatorics concerning Bernoulli trials and the binomial distribution. Poisson distribution is the special case of the binomial distribution when Δ*t* → 0 [51]. The steps of transcription and translation are mechanistically similar and hence, the urn scheme for the Poisson process applies to both. The stationary state count of proteins in a cell is given by a sum of random variables denoting the number of translations per mRNA molecule that is produced in the time needed to reach stationarity. This results in the negative binomial and Neyman type A distributions for the count depending on whether the noise in transcriptions can be ignored or not, respectively [16, 36, 38, 51].

The models for constitutive expression have been extended to include gene regulation. Peccoud and Ycart studied a gene whose promoter switches between active and inactive states [32]. Shahrezaei and Swain extended the Peccoud-Ycart model by accounting for translation and solved it assuming *d*_0_ ≫ *d*_1_, where *d*_1_ is the rate constant for protein degradation [16]. Numerous generalizations and extensions of these models exist and many have been solved analytically, e.g., the leaky two-state model where the promoter switches between two states with different levels of activity [22–24], multi-state models that consider a promoter with more than two states [27, 31, 34, 35, 52], models with auto-regulation [25, 26, 30, 33], etc. All of these models result in probability generating functions that are related to the Kemp families of distributions, which have been derived using various urn models of contagion and population heterogeneity, and as compound or mixture distributions [53–55]. Notably, in each of their applications, the urn model has a distinct design, which is systematically developed to capture the physical characteristics of the natural system under study. Their distinctive features provide an intuitive mapping to the mechanisms behind their respective systems. These urn models have proven fundamental to studies of their intended systems, have immense pedagogical value and have been called a “standard expression” in statistical language [56–58]. For the system of regulated gene expression, while an approach of solving chemical master equations can provide analytical solutions for probability distributions, physical insight into the solutions can be greatly facilitated by the application of urn models. Yet, to the best of our knowledge, urn schemes with well-defined mapping to models of *regulated* gene expression are still lacking.

In this article, we develop an urn model to address this gap and demonstrate its utility by applying it to solve diverse models of gene regulation for stationary state probability distributions. Central to our approach are two principles. First, while transcriptions and translations are regulated by promoter state transitions, arrivals of RNA polymerases or ribosomes occur with fixed propensities independent of the transitions, and can be modeled as unregulated processes. In other words, gene regulation happens because the promoter state determines whether these arrivals result in gene expression, but none of the promoter states exclude polymerases or ribosomes from arriving (Fig. 1a). Second, while there is heterogeneity in promoter activity over long time intervals, i.e., a promoter switches between active and inactive states, in short intervals, the activity is homogeneous. Hence, we map each kinetic model that we consider to an urn model with a master urn (cytosol) and a set of recipient urns of two or more colors (time intervals; see Fig. 1b,c). Each trial in our urn scheme consists of two steps. By virtue of the first principle, the first step of sampling balls (RNA polymerases or ribosomes) from the master urn is done independently of considerations for promoter state transitions. By utilizing the second principle, we devise recipient urns such that each of them represents a time interval with homogeneous promoter activity. In the second step, we assign the balls sampled from the master urn to the recipient urns. We show that the probability distributions of counts of the mRNA and protein molecules from a broad range of models are identical to that of the balls in the recipient urns of a specific color, say red, which represents the active time intervals (Fig. 1d). If the sampling distribution of balls from the master urn is a Poisson distribution and there are recipient urns of two colors, our urn scheme yields the solution of Peccoud and Ycart. If the sampling distribution is negative binomial instead, the urn scheme yields the solution of Shahrezaei and Swain. If there are urns of more than two colors, it yields the solutions for models with multiple promoter states. In all cases, our urn model yields intuitive solutions for the probability distributions and physical interpretations of their parameters with implications for analysis and interpretation of single-cell data.

**Figure 1.**
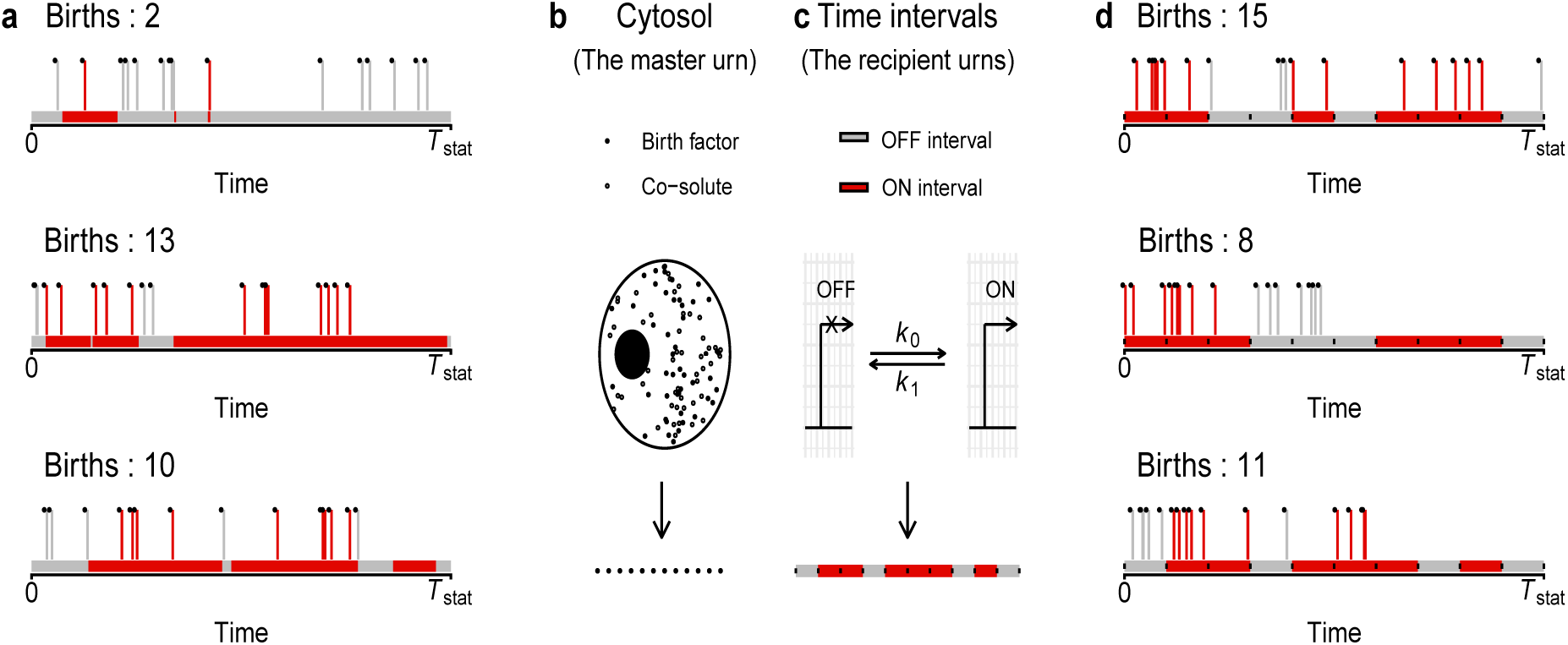
The urn scheme for a system with active and inactive states. **(a)** Three sample trajectories of a gene expression system. The black balls at the heads of spikes are birth factors, which represent RNA polymerases for models of mRNA counts and ribosomes for models of protein counts. The colored bar along the *x*-axis gives the state of the promoter at different time points, with the red color indicating the active state, grey color indicating the inactive state and *T*_stat_ denoting the time scale for stationarity. Births are observed if birth factors arrive when the promoter is active. **(b)** The master urn represents the cytosol of a cell and contains black and white balls, which represent birth factors and solutes other than the birth factors, respectively. Each trial consists of a random sampling of black balls from this urn. **(c)** The recipient urns of red and grey colors represent time intervals with the promoter in the active (ON) or inactive (OFF) states, respectively. Each trial consists of a random ordering of these urns as shown at the bottom. The boundary of each recipient urn is marked with a black dot. **(d)** A trial concludes with a random assignment of the black balls sampled from the master urn to the recipient urns. Assignments to red recipient urns are counted as births.

## Results

### The urn scheme

#### Sampling from the master urn

The master urn represents the cytosol and contains balls of two colors — black and white (Fig. 1b). Each black ball represents a *birth factor*, which we define to be the RNA polymerase for mRNAs and the ribosome for proteins. The white balls represent solutes other than the birth factor. From this urn, we sample black balls over one mean lifetime of the mRNA or protein depending on the time scale, *T*_stat_ for the intended solution to reach stationarity. The sampling process is defined by the kinetic process under consideration. Most models of transcription implicitly assume that the propensity for RNA polymerases to collide with the promoter site does not vary with time but whether a colliding polymerase successfully binds the promoter and transcribes the gene depends on the promoter state. Hence, the number of arrivals follows the Poisson distribution, say with mean *µ* per mRNA lifetime and the urn scheme for Poisson process applies for sampling balls that represent RNA polymerases. We denote the probability distribution of the count, *m*_1_ of polymerase arrivals in one mRNA lifetime as Pois 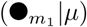, which also denotes the sampling distribution of black balls from the master urn for mRNA distributions (see Supplementary Section 1.1 for a derivation of the Poisson distribution). On the other hand, most models of translation assume that an mRNA molecule is degraded much faster than its protein counterparts, and that in its negligibly short lifetime, it binds a geometrically distributed number of ribosomes resulting in a *burst* of proteins. In our urn scheme, we draw a geometrically distributed number of ribosomes for each potential event of transcription. Hence, the cumulative count, *m*_2_ of ribosome arrivals on mRNAs over one protein lifetime follows the negative binomial distribution with parameters *α* and *β*, which represent the number of potential transcriptions in one protein lifetime and the mean number of ribosomes that bind to each mRNA, respectively (the interpretation of *α* is given in more detail later). We denote this NB 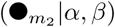 (see Supplementary Section 1.2).

### Assignment to the recipient urns

Each recipient urn represents a time interval with a uniform propensity for gene expression. Each urn has one of two or more colors, with the number of colors dependent on the number of promoter states (Fig. 1c). We defer the mathematical exposition of our urn scheme to a later section. Here, it suffices that at stationary state, the time intervals captured by urns are equal and represent the characteristic time scale of promoter state transitions. Furthermore, the total time captured by the set of recipient urns equals *T*_stat_. For a genetic switch with two states, let grey and red urns represent the inactive and active states, respectively. Also, say *n*_grey_ and *n*_red_ represent their numbers, respectively. At stationary state, *n*_red_ and *n*_grey_ are fixed and their permutations represent samples of state-transition trajectories. The interaction of the time points of arrivals of birth factors with the promoter state transitions determines the outcome of interest, which is the number of births (Fig. 1d). In the urn model parlance, we capture this by assigning the black balls from the master urn to the recipient urns. If a ball is assigned to a red urn, a birth is observed. The outcome of interest is the number of balls assigned to the red urns collectively. Since the times of arrivals of birth factors are independent of the trajectory of promoter state transitions, the process of assigning balls to the recipient urns allows equal likelihood of assignment to all urns. Let us say that in an experiment, we sample *m* + *i* black balls from the master urn. The probability that a random assignment to the recipient urns results in exactly *m* balls assigned to the red urns is given by a negative hypergeometric distribution. To see this, note that the number of ways to divide *m* + *i* balls in *n*_red_ + *n*_grey_ urns is 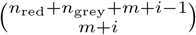, which is the same as the number of ways to permute *m* + *i* identical balls and *n*_red_ + *n*_grey_ − 1 identical dividers [51]. Of these, 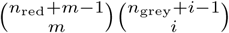 are such that exactly *m* balls are assigned to the red urns. Hence, the probability of the outcome *m* is the ratio of this quantity to the total number of ways to assign *m* + *i* balls and we denote the distribution as NH (●_*m*_ ↦ *n*_red_ | ●_*m*+*i*_ ↦ {*n*_red_, *n*_grey_}), where ‘↦’ represents the process of assignment (see Supplementary Section 2).

### Application to the Peccoud-Ycart model

The Peccoud-Ycart model for probability distribution of mRNA counts considers a promoter that can exist in active and inactive states (Fig. 2a). Using the urn model parlance here, the black balls represent RNA polymerases. To observe an outcome of *m*_1_ transcriptions, the sample drawn from the master urn must contain *m*_1_ or more black balls, say *m*_1_ + *i*_1_ with *i*_1_ ≥ 0. The probability of this event is Pois 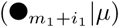. Given *m*_1_ + *i*_1_ balls, *n*_red_ red recipient urns and *n*_grey_ grey recipient urns, the probability that exactly *m*_1_ balls are assigned to the red urns is NH 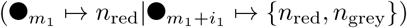. The joint probability of *m*_1_ balls assigned to the red urns and *i*_1_ to the grey urns is given by the product of the said Poisson and negative hypergeometric distributions. A marginal of the joint probability yields the probability of *m*_1_ transcriptions,

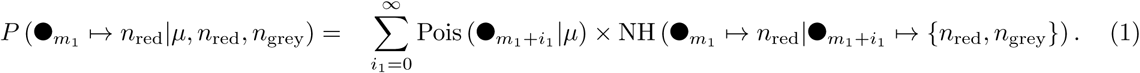

**Figure 2.**
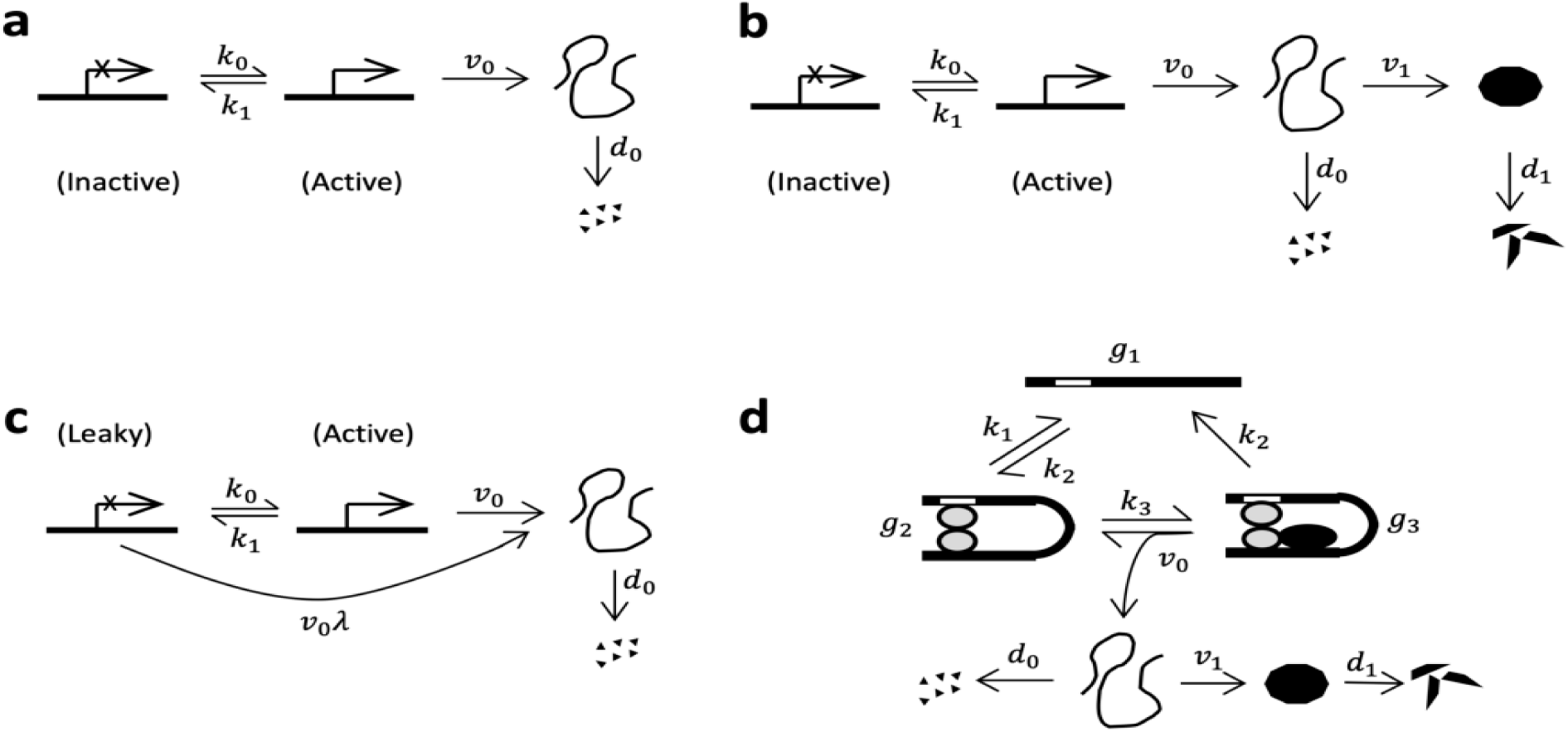
Kinetic schemes for regulated gene expression. **(a)** The Peccoud-Ycart model considers a promoter that switches between inactive and active states with propensities *k*_0_ and *k*_1_. The active state transcribes with propensity *v*_0_ and mRNAs are degraded with propensity *d*_0_. **(b)** The Shahrezaei-Swain model builds on the Peccoud-Ycart model by accounting for translations of mRNAs with propensity *v*_1_ and degradation of proteins with propensity *d*_1_. **(c)** The leaky two-state model generalizes the Peccoud-Ycart model by replacing the inactive state with a leaky state that transcribes with propensity *v*_0_*λ*. **(d)** Cao *et al.* consider a promoter that transitions between three states — *g*_1_, *g*_2_ and *g*_3_. In state *g*_1_, neither the transcription factor nor the RNA polymerase are bound to the promoter. In state *g*_2_, the transcription factor is bound to the promoter and in state *g*_3_, both the polymerase and the transcription factor are bound. State *g*_3_ releases the polymerase with propensity *v*_0_, which results in transcription and translations as well as a transition to *g*_2_. In addition, the model allows reversible transitions between *g*_1_ and *g*_2_ with propensities *k*_1_ and *k*_2_, transition from *g*_2_ to *g*_3_ with propensity *k*_3_ and transition from *g*_3_ to *g*_1_ with propensity *k*_2_.

Its generating function is given in terms of the Kummer’s hypergeometric function of the first kind, 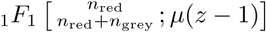, which is formally identical to the solution of Peccoud and Ycart (see Supplementary Section 3.1 for the proof). We defer the mapping of the kinetic parameters to parameters of the urn model to a later section.

### Application to the Shahrezaei-Swain model

Similarly to the Peccoud-Ycart model, the Shahrezaei-Swain model considers transcriptionally inactive and active states but also accounts for translation to solve for the probability distribution of protein counts (Fig. 2b). Hence, for urn modeling in this case, we let the black balls represent ribosomes. To observe *m*_2_ translations, the sample drawn from the master urn must contain *m*_2_ + *i*_2_ black balls with *i*_2_ ≥ 0, which happens with probability NB 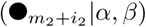. Once again, we consider *n*_red_ red recipient urns and *n*_grey_ grey recipient urns, but in this case they represent *translationally* active or inactive intervals, respectively. Note that a subset of recipient urns that represented transcriptionally active time intervals in the case of Peccoud-Ycart model might not receive any polymerase arrivals, which renders them translationally inactive. Hence, *n*_red_ and *n*_grey_ have a different mapping to the kinetic parameters than in case of the Peccoud-Ycart model, as we show later. Regardless, given *n*_red_ and *n*_grey_, the probability of *m*_2_ translations follows from similar arguments as before,

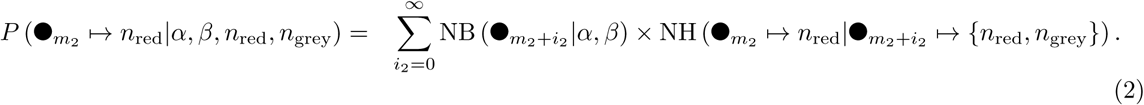

The generating function for this distribution is given in terms of the Gaussian hypergeometric function, 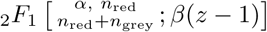, which is formally identical to the solution of Shahrezaei and Swain (see Supplementary Section 3.2 for the proof).

### Application to the leaky two-state model

The leaky two-state model for mRNAs considers two states with propensities of transcription differing by a constant factor, say *λ* (Fig. 2c). Let the expected numbers of arrivals of RNA polymerase in one mRNA lifetime be *µλ* and *µ* in the leaky and fully active states, respectively, with 0 < *λ* < 1. Since we consider a model for probability distribution of mRNA counts, the black balls drawn from the master urn represent RNA polymerases. In this case, we consider two parallel experiments in our urn scheme, which represent the urn with the contributions of a constitutive component with the expected value of *µλ* and a regulated component with the expected value of *µ* (1 − *λ*). In the first experiment, we draw a sample of *m*_1,*a*_ black balls from the master urn with the probability Pois 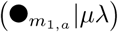, which represents the leakage that is unaffected by promoter state transitions. Hence, with respect to this sample, all recipient urns are red and all of it is counted towards transcriptions. In the second experiment, we draw a sample of *m*_1,*b*_ + *i*_1_ balls from the master urn with the probability Pois 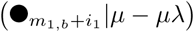, which represents the regulated component of polymerase arrivals. This experiment proceeds as described earlier for the Peccoud-Ycart model and allows us to derive the probability of *m*_1,*b*_ transcriptions from the regulated component by substituting *m*_1_ with *m*_1,*b*_ and *µ* with *µ* (1 − *λ*) in Eq. 1. The overall outcome of interest is *m*_1_ = *m*_1,*a*_ + *m*_1,*b*_. Its probability is given by the convolution rule and has a generating function that is the product of the generating functions for Pois 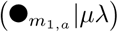 and that for the regulated component, i.e., 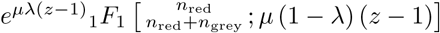, which is consistent with the solution of Cao and Grima [22]. Note that previously, we have shown in the context of a model for the *lac* operon of *E. coli* that this generating function can be viewed as a convolution of contributions from a leaky sub-system of Lac repressor-bound states and regulated transitions to the repressor-free state [24]. A recent preprint also utilizes an identical concept [59]. In this section, we have shown that the same concept is easily accommodated in the framework of our urn model. Further, the distribution of proteins from a leaky two-state model can be derived similarly.

### Application to models with multiple states

Analytical solutions are available for models with more than two but a fixed number of promoter states [27, 34] as well as those with an arbitrary number of states [31]. Next, we consider the model by Cao *et al.*, where the promoter exists in three states, say *g*_1_, *g*_2_ and *g*_3_ with *g*_3_ being active (Fig. 2d). For the mRNA counts, the sampling distribution of *m*_2_ + *i*_2_ + *j*_2_ balls (RNA polymerases) from the master urn is Pois 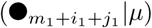, where *i*_2_, *j*_2_ ≥ 0. We account for the additional promoter state by adding another layer of recipient urns. First, we divide the balls between urns that correspond to the sub-system of state *g*_1_ and the sub-system of *g*_2_ and *g*_3_ collectively. Let there be *n*_blue_ blue and *n*_ppl_ purple urns for these sub-systems, respectively. Then, the probability of assigning *m*_2_ + *i*_2_ balls to the purple urns is NH 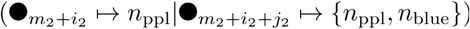. The balls assigned to the purple urns are further re-assigned to another layer of red and grey recipient urns — *n*_red_ and *n*_grey_ in number, respectively. Let the grey and red urns represent the states *g*_2_ and *g*_3_ respectively. The outcome of interest, assignment of *m*_2_ balls to the red recipient urns has the probability NH 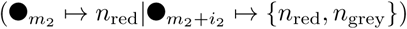. Hence, the probability of *m*_2_ transcriptions is

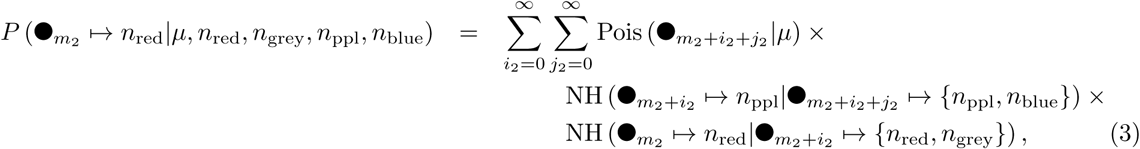

which has a generalized hypergeometric function, 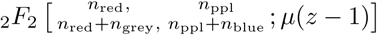 as its generating function (see Supplementary Section 3.3 for the proof). This solution can be extended to the protein distribution for a model with two inactive and one active promoter states by replacing the Poisson distribution with the negative binomial distribution. This yields the solution by Cao et al. [27] (see Supplementary Section 3.4 for the proof). It can be extended to a model of an arbitrary number of promoter states with all but one inactive by adding additional layers of recipient urns. Then, if sampling from the master urn follows the Poisson distribution, we get the solution by Zhou and Liu for the probability distribution of mRNA counts [31] (see Supplementary Section 3.5 for the proof). Once again, if sampling from the master urn follows the negative binomial distribution, we get the solution for protein counts (see Supplementary Section 3.6).

### Relationship between the kinetic and urn model parameters

We have shown that our urn model yields probability distributions that are formally identical to those from kinetic models. In this section, we obtain the relationship between parameters of the kinetic and urn models. To this end, say, we follow a cell in real time starting at time *t* = 0 when the gene system under consideration is at stationary state (e.g., Fig. 1a shows trajectories of three cells). We define that event *E*_*m*_ occurs when *m* births are observed in a time interval given by the time scale, *T*_stat_ for reaching stationarity (e.g., the top panel in Fig. 1a illustrates the event *E*_2_). Let *M* be a random variable representing the number of arrivals of birth factors in *T*_stat_ time (the number of black balls in any trajectory shown in Fig. 1a), {*T*_1_, *T*_2_, …, *T*_*M*_} be the random variables representing the time points of arrivals (*x*-coordinates of the spikes in Fig. 1a), and *R*_ON_ be a random variable representing the set of time points when the promoter is active (set of all the time points in red colored segments along the *x*-axis in any trajectory shown in Fig. 1a). Then, *E*_*m*_ occurs when 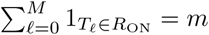, where 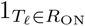 is an indicator variable that equals 1 if *T*_𝓁_ ∈ *R*_ON_ and 0 otherwise. For this to happen, there must be at least *m* arrivals of the birth factors in total, i.e., *M* = *m* + *i* such that *i* ≥ 0. Given that this condition is met, there must be exactly *m* arrivals during the active time intervals, i.e.,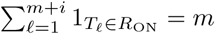. Essentially, we find that the event of *m* births can be decomposed into two simpler events, whose probabilities can be derived separately. Next, let us define *R*_red_ as the counterpart of *R*_ON_ in the urn space (Fig. 1d). In other words, *R*_red_ is the subset of time points from the set [0, *T*_stat_] that fall in red urns for a sample permutation of the recipient urns. Say, *E*_urn,*m*_ represents the event 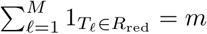. Then, we must choose *n*_red_ and *n*_grey_ such that *P* (*E*_*m*_) = *P* (*E*_urn,*m*_).

For the Peccoud-Ycart model, 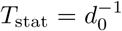 and RNA polymerases function as the birth factors. As we mentioned, the sampling of polymerases from the master urn can be modeled as a Poisson process, which yields 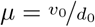. Next, we solve for *n*_red_ and *n*_grey_. Since the *T*_𝓁_’s are independent of each other, 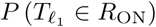 is independent of 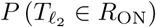 for 𝓁_1_ ≠ 𝓁_2_. Hence, *P* (*E*_*m*_) = *P* (*E*_urn,*m*_) if *P* (*T*_𝓁_ ∈ *R*_ON_) = *P* (*T*_𝓁_ ∈ *R*_red_) for all 𝓁. Let the propensities for promoter state transitions be *k*_0_ and *k*_1_ (Fig. 2a). At stationary state, 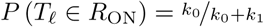. To derive *P* (*T*_𝓁_ ∈ *R*_red_), let *w*_red_ and *w*_grey_ represent the time duration captured by each of the red and grey urns, respectively, and min (*w*_red_, *w*_grey_) be the minimum of the two. Then, for *T*_𝓁_ close to the boundaries of the set [0, *T*_stat_], i.e. for *T*_𝓁_ less than min (*w*_red_, *w*_grey_) away from the boundaries, 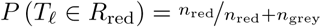. This is because all of the duration from *t* = 0 to min (*w*_red_, *w*_grey_) is contained within a single urn, which can either be red or grey. On the other hand, for an arbitrary choice of *T*_𝓁_, 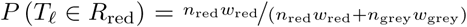. In other words, the probability that a randomly chosen time point falls in a red urn is given by the fraction of time in the interval [0, *T*_stat_] that is covered by the red urns. At stationary state, whether *T*_𝓁_ falls in *R*_red_ should be the same for all *T*_𝓁_, which requires *w*_red_ = *w*_grey_. Let us replace *w*_red_, *w*_grey_ with *w*. Now, if we were to arbitrarily pick a polymerase arrival and shift the corresponding *T*_𝓁_ to the left or right by a fixed amount (say, by grabbing one of the spikes in Fig. 1d), whether the shifted spike still falls in an urn of the same color depends on *w*. In simple words, while there is temporal heterogeneity in transcriptional activity over long periods, in short time windows around any event of polymerase arrival, transcriptional activity is homogeneous. Given a suitable choice of *w*, these time windows can be modeled as if they were composed of urns of the same color. *w* is determined by the transient time scale for switching of transcriptional activity, which is *w* = (*k*_0_ + *k*_1_)^−1^ (see the section on reversible unimolecular reactions in McQuarrie [60] for derivation of the transient time scale). Finally, for *P* (*T*_𝓁_ ∈ *R*_ON_) = *P* (*T*_𝓁_ ∈ *R*_red_), 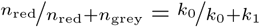. Since, 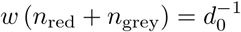, we obtain 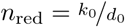 and 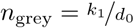. Essentially, we find that *n*_red_ and *n*_grey_ are equal to the normalized propensities for the promoter to switch to the active and inactive states, respectively. By replacing these parameter values in Eq. 1, we retrieve the analytical solution of Peccoud and Ycart.

Shahrezaei and Swain solved a kinetic model for protein production assuming *d*_0_ ≫ *d*_1_, which implies 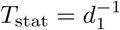. In this case, ribosomes function as the birth factors and for the system to be active for protein production, the gene must be transcriptionally active and RNA polymerase arrivals must occur. In other words, if we start our observation with the promoter in the transcriptionally inactive state, the waiting time for transition to the translationally active state is greater than 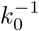. In the remaining part of this section, we first derive the propensity for translational activity, and then apply the approach described above to obtain *n*_red_ and *n*_grey_ for the Shahrezaei-Swain model. To this end, let us pick a time point randomly as *t* = 0. Next, we define *p*_0_ (*t*) as the probability that the gene is transcriptionally inactive at time *t* and no mRNA has been produced in time [0, *t*]. Similarly, *p*_1_ (*t*) is the probability that the gene is transcriptionally active at time *t* but no mRNA has been produced in time [0, *t*]. Then,

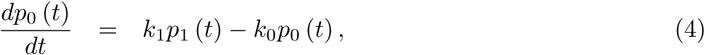

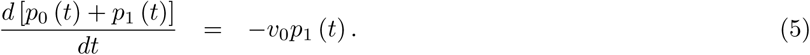

We are interested in the time scale at which *p* (*t*) = *p*_0_ (*t*) + *p*_1_ (*t*) approaches 0, i.e. the time scale at which the marginal probability of no mRNA production decays to 0. To this end, Eqs. 4-5 can be combined to get a second order differential equation,

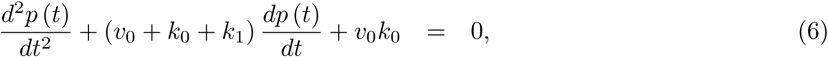

which admits solutions of the form 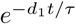, where *τ* > 0 is a characteristic time scale of the system. Substituting 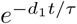 in Eq. 6 yields a quadratic equation, with the roots

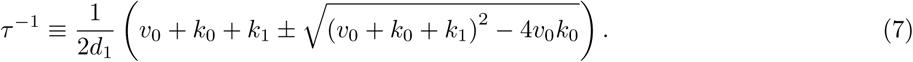

Inverse of the larger of the roots, i.e. the slow time scale (say, *τ*_*s*_) gives the waiting time for mRNA production and the inverse of the smaller one yields the fast time scale (say, *τ*_*f*_). We interpret 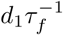 as the propensity for occurrence of any event, i.e., promoter state transition or polymerase arrival. For example, if we know that a polymerase arrival occurs at any time point *t*, it is likely that there will be no arrival in the time window 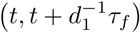 because the polymerase arrival occurs as fast as the fast time scale of the system allows. Now, we can derive the relationship between the kinetic and urn model parameters in terms of these time scales. First, we sample ribosomes from the master urn, such that for each 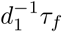 time window in *T*_stat_, we draw a geometrically distributed number of ribosomes. The geometric distribution has the mean 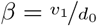 and we draw 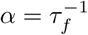 geometrically distributed samples. Hence, the probability distribution for a sample of *m*_2_ + *i*_2_ ribosomes is given by the negative binomial distribution with the said values for *α* and *β*. Next, we distribute these in the red and grey recipient urns. Similarly to arrivals of polymerases, arrivals of ribosomes are independent of each other. We can use the same method as described for Peccoud-Ycart model to derive *n*_red_ and *n*_grey_. The difference is that here, red urns represent translationally active time windows. Hence, their number is given by 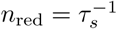 and 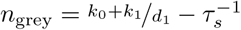, which are the normalized propensities for the system to switch to the translationally active and inactive states, respectively. By using these parameter values in Eq. 2, we retrieve the solution of Shahrezaei and Swain.

### Solving the urn model using the inclusion-exclusion principle

In the previous sections, we have written the probability of *exactly m*_1_ assignments to the red urns out of a sample of *m*_1_ + *i*_1_ balls from the master urn directly as a negative hypergeometric distribution. The same probability could be written from an alternative perspective, where we start with the probabilities for at least *i*_1_ assignments to the grey urns, at least *i*_1_ + 1 assignments to the grey urns, …, at least *i*_1_ + *m*_1_ − 1 assignments to the grey urns and all *i*_1_ + *m*_1_ assignments to the grey urns. Then, we combine these as an alternating sum using the principle of inclusion-exclusion to write the probability of exactly *m*_1_ assignments to the red urns (see Supplementary Section 4). For the Peccoud-Ycart model, this approach yields an expression for 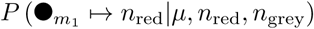 that is the 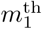 coefficient in the Maclaurin series expansion of Kummer transformation of the Peccoud-Ycart solution, 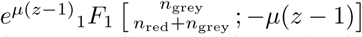 with 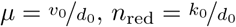 and 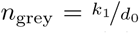. This way of representing the Peccoud-Ycart solution leads to an interpretation of the transcriptional dynamics in terms of transcriptional *lapses* as we show next, and contrast with the dynamics of transcriptional bursts.

For a repressed gene with 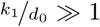, the hypergeometric function, 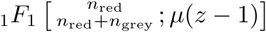 reduces to the generating function for the negative binomial distribution, 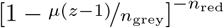 [61]. Here, 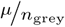 is interpreted as the mean number of mRNAs produced when the promoter switches to the active state (burst size), *n*_red_ as the frequency of switching to the active state (burst frequency), and the hypergeometric function is the generating function for the Peccoud-Ycart solution. In Supplementary Section 5.1, we use a perturbation theoretic approach to show that this is valid for *k*_1_ ≫ *k*_0_. U sing this limiting form of the hypergeometric function, when *k*_0_ ≫ *k*_1_, i.e., for an activated gene, 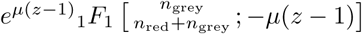 reduces to 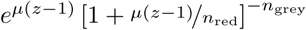 (see Supplementary Section 5.2 for proof). Hence, in both the activated and repressed scenarios, the limiting form of the Peccoud-Ycart solution has a term of the kind [1 − *β* (*z* − 1)]^−*α*^, which is the generating function for a negative binomial distribution if *α, β* > 0. However, 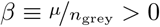 in the repressed limit but 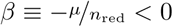 in the activated limit. Hence, this term is not consistent with the interpretation of transcriptional dynamics in the activated case as being bursty and 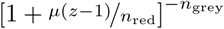 is not a probability distribution. We interpret the dynamics in the activated case as one that is composed of transcriptional lapses, whereby the gene is expressed with uniform propensity *v*_0_ for majority of the time and for short-lived intervals, there is a lapse in transcriptional activity brought about by gene regulation (Fig. 3a). In Fig. 3b, we show that for a gene in the activated case, the approximation in terms of transcriptional lapses is in significantly better agreement with the exact solution than that in terms of transcriptional bursts. This approximation might be useful in fitting single-cell data for activated genes.

**Figure 3.**
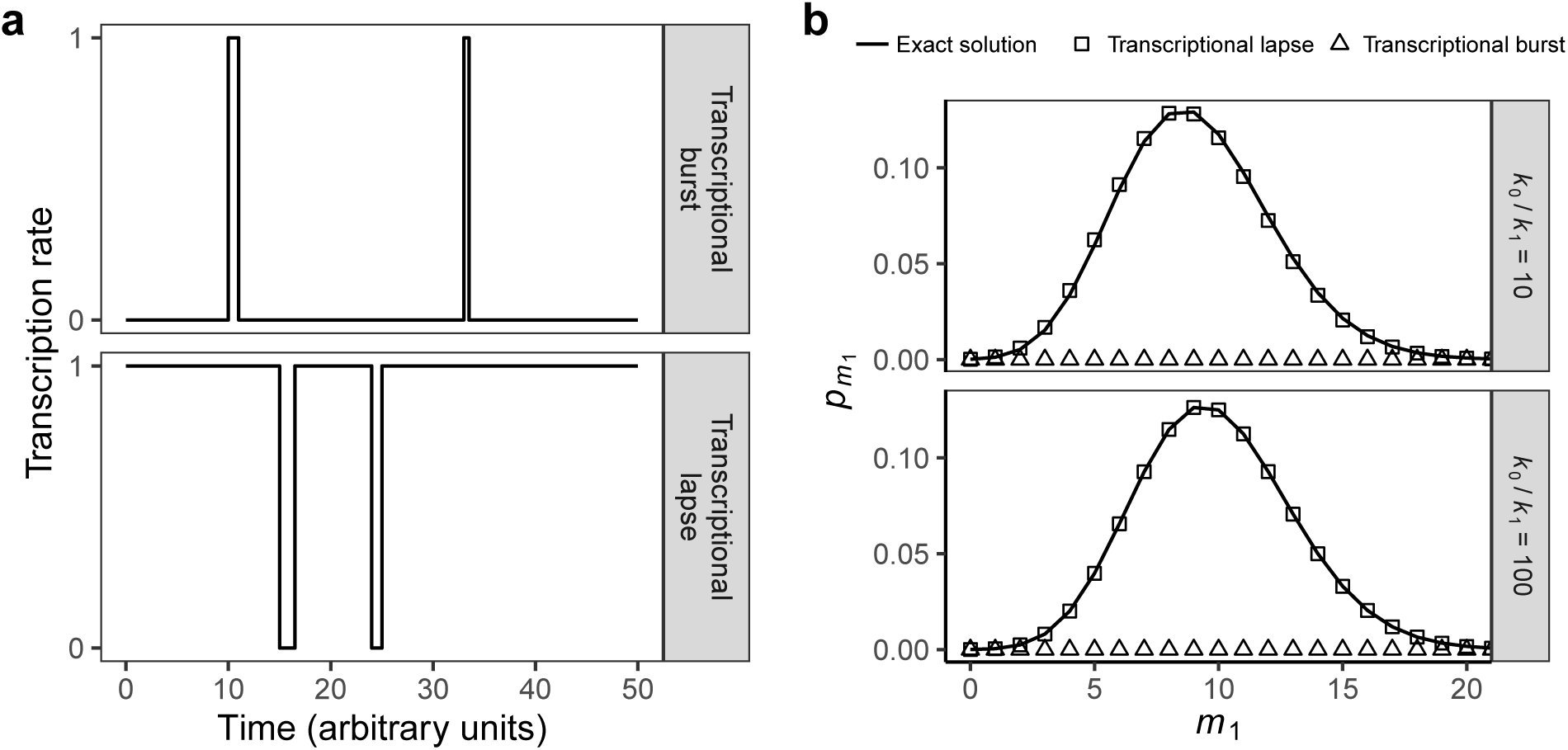
The dynamics of transcription for activated genes involve transcriptional lapses. **(a)** The transcriptional dynamics of repressed genes are characterized by short-lived periods of activity (bursts) and that of activated genes by short-lived periods of inactivity (lapses). **(b)** Stationary state probability of *m*_1_ mRNAs obtained from exact solution of the Peccoud-Ycart model (solid line) and its limiting forms for transcriptional lapses (rectangles) and bursts (triangles). Both the panels show the probabilities in the activated case, when *k*_0_ is 10 times *k*_1_ (top) or 100 times *k*_1_ (bottom) with *k*_1_ = 0.01 s^−1^, *v*_0_ = 0.05 s^−1^ and *d*_0_ = 0.005 s^−1^. The peaks from the transcriptional burst model appear at *m*_1_ ≫ 10 for both the panels.

## Discussion

### Comparison with existing models for the Kemp families of distributions

All problems concerning probabilities could be addressed using urn models [57]. However, it is a non-trivial task to develop physically meaningful urn schemes for any given system, even if solutions for the probability distributions of interest were known. This is because many different models can give rise to identical distributions [56]. Indeed, this is true for distributions resulting from models of regulated gene expression, which are related to the Kemp families of distributions that are well-known in the urn modeling literature [62]. A general feature of some of the models that lead to such distributions is that in a series of trials, multiple instances of the same outcome tend to occur in close succession. This could result due to the presence of contagion or population heterogeneity in the stochastic process under consideration [54]. We consider a known model for each and discuss how they differ from models of regulated gene expression. The Pólya-Eggenberger urn model studies the spread of contagious diseases [56]. In this model, one considers a finite urn with a pre-specified number of white and black balls, and draws balls from the urn randomly and one at a time. The drawn ball is replaced along with additional balls of the same color, which increases the probability of drawing this color in the next trial. This process simulates the spread of pathogens once they appear in a population. However, in the case of regulated gene expression, such contagion does not exist because the concentrations of birth factors do not change with time. Hence, the process of replacing a ball with additional balls of the same color is not physically meaningful. Alternatively, Gurland considered a compound Poisson process, which is a Poisson process with a rate parameter that is also a random variable due to heterogeneity of the population in consideration [54]. For example, consider a constitutively expressed gene in a population of cells such that the rate of RNA polymerase arrival at the promoter varies from cell to cell. If the rate parameter depended on a random variable with Beta distribution, the probability distribution of the corresponding mRNA counts would be identical to the solution of the Peccoud-Ycart model. It is noteworthy that this interpretation has been recently utilized to fit single-cell RNA-seq data [45, 46]. However, the Peccoud-Ycart model, like most mechanistic models of gene expression, is solved for a homogeneous population with a fixed rate of polymerase arrivals. Hence, the interpretation as a compound Poisson process is not consistent with its kinetic scheme.

### Implications for analysis of single-cell data

Our study provides helpful insights for single-cell data analysis. First, we find that the urn model parameters are related to the time scales of switching between sub-systems of the promoter states. Particularly, in the case of the Shahrezaei-Swain model, the numerator parameters of the Gaussian hypergeometric generating function are related to the fast time scale and the waiting time for transcriptions. To the best of our knowledge, these parameters have not been ascribed physical interpretations previously. Our interpretations would facilitate mechanistic understanding from analysis of single-cell data for proteins if the data were fit using the Shahrezaei-Swain solution. Second, the field has recently witnessed a rapid development of single-cell technologies and parallel advances in analytical solutions of models incorporating mechanistic details of gene expression [27, 63, 64]. As such, solutions of the detailed models would likely be utilized for analyzing single-cell data in the future. We find that solutions of disparate mechanistic models (e.g., the models of Cao et al. and Karmakar [27, 34]) may result in identical distributions if they involve the same number of timescales for switching between the sub-systems of promoter states. Hence, if a probability distribution yields good fits to data on mRNA or protein counts, it indicates that the number of time scales represented in the distribution is adequate to describe the gene system. Additional data must be collected to distinguish between the mechanistic models leading to these time scales as well as to assess whether the probability distribution might have resulted from the presence of features such as contagion or population heterogeneity in the system. Further insights in this direction could be gained from existing statistical literature to distinguish between such features [65]. Finally, we note that in some cases, the researchers utilize the negative binomial distribution to fit their data [11]. This could be motivated by the challenges of computing hypergeometric functions for the complete range of parameter values [66]. Furthermore, the parameters of negative binomial distribution have well-understood interpretations as the burst size and frequency of expression. We find that at least for activated genes, the transcriptional dynamics are better described in terms of transcriptional lapses. In the general case, it might be worthwhile to consider a distribution with the generating function *e*^*µ*(*z*−1)^ [1 − *β* (*z* − 1)]^−*α*^ to fit the data. If all the parameters, *α, β* and *µ* were non-negative, this is a Delaporte distribution, which could be interpreted in terms of a leaky two-state model with transcriptional bursts (see Supplementary Section 5) [62]. If *β* were allowed to be negative, the parameters could be interpreted in terms of a leaky two-state model with transcriptional lapses.

## Limitations

Limitations of our approach are worth considering. First, as the mechanistic models grow in complexity, identifying the relationship between the urn model and kinetic parameters becomes a challenging task. Despite this limitation, our approach complements the existing approaches for solving the stochastic models by revealing the origins of the solutions in terms of the physics underlying the dynamics of gene regulation. Second, our model assumes that birth factors arrive independently of each other and that there are no feedback loops. If these assumptions do not hold, the birth factors cannot be assigned to the recipient urns independently of each other and the numbers of recipient urns of different colors are not independent of the counts of births. Nevertheless, the solution for a model with feedback could be viewed as correction terms combined with the solution for its simplified version that ignores feedback. Further, we note that, like most models that have been solved analytically, our model implicitly assumes that the reactants and catalysts needed for gene expression such as tRNAs, amino acids, ribonucleotides, etc. are available in sufficient quantity [5]. In the future, generalizations of our model might be able to relax these assumptions. Third, our model makes physical sense only if the parameters such as *n*_red_ and *n*_grey_ are integer-valued. This might not be true for arbitrary values of the kinetic parameters. In this case, the factorial functions appearing in our expressions must be replaced by gamma functions as done for the urn model of the negative binomial distribution [16].

## Conclusion

We developed an urn model and applied it to study a broad range of stochastic models of regulated gene expression. Our urn model generalizes the classical birth-death model by considering the regulation of births. The classical model makes no distinction between arrivals of birth factors and births, as all arrivals cause births. However, in the presence of regulation, as in the case of gene expression, arrivals of birth factors do not result in births if they occur when genes are inactive. Hence, we reinterpret the classical scheme of births as a scheme to sample birth factors from a master urn, which represents the cytosol of a cell. Next, we note that despite temporal heterogeneity in expression activity of genes over long time intervals, there is homogeneity in short intervals. We use this concept to devise recipient urns of two or more colors, which are discretized time intervals with the promoter existing in a single sub-system of its states for the duration of each urn. Then, we assign the birth factors to the recipient urns and count the assignments to urns of a specific color as births. Given physically intuitive choices of sampling distribution from the master urn and the numbers of recipient urns for each color, our model yields probability distributions that are identical to solutions of a range of kinetic models. We describe the physical principles that lead to our urn scheme and provide kinetic interpretations for the urn model parameters. Finally, we discuss our approach in the broader context of urn models and single-cell data analysis as well as highlight its limitations. We conclude by noting that the solutions from chemical master equations are obtained in terms of generating functions and physical intuition into origins of the expressions for probability distributions have thus far been limited for models with regulation. Our model facilitates direct interpretation of the probability expressions, which underscores its significance for pedagogical purposes and to interpret results from data fitting. The physical insights developed in this work will facilitate the adoption of the analytical solutions in single-cell studies.

## Supporting Information

Supplementary information for this article is available online.

## Author contributions

KC and AN conceived the project. KC designed the research, developed the method, generated the figures and wrote the manuscript. AN provided critical feedback and helped shape the research, analysis and manuscript. KC did the proofs other than the perturbation theoretic proofs, which were done by AN. Both the authors read and approved the manuscript.

## References

1. George Orphanides and Danny Reinberg. A unified theory of gene expression. Cell, 108(4):439–451, 2002.

2. Arjun Raj and Alexander van Oudenaarden. Nature, nurture, or chance: stochastic gene expression and its consequences. Cell, 135(2):216–226, 2008.

3. Avigdor Eldar and Michael B Elowitz. Functional roles for noise in genetic circuits. Nature, 467(7312):167, 2010.

4. Mads Kaern, Timothy C Elston, William J Blake, and James J Collins. Stochasticity in gene expression: from theories to phenotypes. Nature Reviews Genetics, 6(6):451, 2005.

5. Johan Paulsson. Models of stochastic gene expression. Physics of life reviews, 2(2):157–175, 2005.

6. Roy D Dar, Brandon S Razooky, Abhyudai Singh, Thomas V Trimeloni, James M McCollum, Chris D Cox, Michael L Simpson, and Leor S Weinberger. Transcriptional burst frequency and burst size are equally modulated across the human genome. Proceedings of the National Academy of Sciences, 109(43):17454–17459, 2012.

7. Caroline R Bartman, Nicole Hamagami, Cheryl A Keller, Belinda Giardine, Ross C Hardison, Gerd A Blobel, and Arjun Raj. Transcriptional burst initiation and polymerase pause release are key control points of transcriptional regulation. Molecular cell, 73(3):519–532, 2019.

8. Shasha Chong, Chongyi Chen, Hao Ge, and X Sunney Xie. Mechanism of transcriptional bursting in bacteria. Cell, 158(2):314–326, 2014.

9. Jonathan M Raser and Erin K O’Shea. Control of stochasticity in eukaryotic gene expression. Science, 304(5678):1811–1814, 2004.

10. Keisuke Fujita, Mitsuhiro Iwaki, and Toshio Yanagida. Transcriptional bursting is intrinsically caused by interplay between RNA polymerases on DNA. Nature communications, 7:13788, 2016.

11. Lok H. So, Anandamohan Ghosh, Chenghang Zong, Leonardo A. Sepulveda, Ronen Segev, and Ido Golding. General properies of the transcriptional time-series in *E. coli*. Nature Genetics, 43(6):554–560, 2011.

12. Daniel L Jones, Robert C Brewster, and Rob Phillips. Promoter architecture dictates cell-to-cell variability in gene expression. Science, 346(6216):1533–1536, 2014.

13. Tineke L Lenstra, Joseph Rodriguez, Huimin Chen, and Daniel R Larson. Transcription dynamics in living cells. Annual review of biophysics, 45:25–47, 2016.

14. Brian Munsky, Gregor Neuert, and Alexander Van Oudenaarden. Using gene expression noise to understand gene regulation. Science, 336(6078):183–187, 2012.

15. KA Geiler-Samerotte, CR Bauer, S Li, N Ziv, David Gresham, and ML Siegal. The details in the distributions: why and how to study phenotypic variability. Current opinion in biotechnology, 24(4):752–759, 2013.

16. Vahid Shahrezaei and Peter S Swain. Analytical distributions for stochastic gene expression. Proceedings of the National Academy of Sciences, 105(45):17256–61, 2008.

17. Alvaro Sanchez, Hernan G. Garcia, Daniel Jones, Rob Phillips, and Jané Kondev. Effect of promoter architecture on the cell-to-cell variability in gene expression. PLoS Computational Biology, 7(3):e1001100, 2011.

18. JM Vilar and L Saiz. Suppression and enhancement of transcriptional noise by DNA looping. Physical review. E, Statistical, nonlinear, and soft matter physics, 89(6):062703–062703, 2014.

19. Tyler M Earnest, Elijah Roberts, Michael Assaf, Karin Dahmen, and Zaida Luthey-Schulten. DNA looping increases the range of bistability in a stochastic model of the *lac* genetic switch. Physical biology, 10(2):026002, 2013.

20. Thomas B Kepler and Timothy C Elston. Stochasticity in transcriptional regulation: origins, consequences, and mathematical representations. Biophysical journal, 81(6):3116–3136, 2001.

21. Masaki Sasai and Peter G Wolynes. Stochastic gene expression as a many-body problem. Proceedings of the National Academy of Sciences, 100(5):2374–2379, 2003.

22. Zhixing Cao and Ramon Grima. Linear mapping approximation of gene regulatory networks with stochastic dynamics. Nature communications, 9(1):3305, 2018.

23. Krishna Choudhary, Stefan Oehler, and Atul Narang. Protein distributions from a stochastic model of the *lac* operon of *E. coli* with DNA looping: Analytical solution and comparison with experiments. PLoS One, 9(7):1–14, 2014.

24. Krishna Choudhary and Atul Narang. Analytical expressions and physics for single-cell mRNA distributions of the *lac* operon of *E. coli*. Biophysical journal, 117(3):572–586, 2019.

25. Ramon Grima, Deena R Schmidt, and Timothy J Newman. Steady-state fluctuations of a genetic feedback loop: An exact solution. The Journal of Chemical Physics, 137(3):035104, 2012.

26. Yves Vandecan and Ralf Blossey. Self-regulatory gene: an exact solution for the gene gate model. Physical Review E, 87(4):042705, 2013.

27. Zhixing Cao, Tatiana Filatova, Diego A Oyarzún, and Ramon Grima. Multi-scale bursting in stochastic gene expression. bioRxiv, page 717199, 2019.

28. Zhixing Cao and Ramon Grima. Analytical distributions for detailed models of stochastic gene expression in eukaryotic cells. Proceedings of the National Academy of Sciences, 2020.

29. Hodjat Pendar, Thierry Platini, and Rahul V Kulkarni. Exact protein distributions for stochastic models of gene expression using partitioning of poisson processes. Physical Review E, 87(4):042720, 2013.

30. José EM Hornos, Daniel Schultz, Guilherme CP Innocentini, JAMW Wang, Aleksandra M Walczak, José N Onuchic, and Peter G Wolynes. Self-regulating gene: an exact solution. Physical Review E, 72(5):051907, 2005.

31. Tianshou Zhou and Tuoqi Liu. Quantitative analysis of gene expression systems. Quantitative Biology, 3(4):168–181, 2015.

32. Jean Peccoud and Bernard Ycart. Markovian modeling of gene product synthesis. Theoretical Population Biology, 48:222–234, 1995.

33. Niraj Kumar, Thierry Platini, and Rahul V Kulkarni. Exact distributions for stochastic gene expression models with bursting and feedback. Physical review letters, 113(26):268105, 2014.

34. Rajesh Karmakar. Conversion of graded to binary response in an activator-repressor system. Physical Review E, 81(2):021905, 2010.

35. Huahai Qiu, Bengong Zhang, and Tianshou Zhou. Influence of complex promoter structure on gene expression. Journal of Systems Science and Complexity, pages 1–15, 2018.

36. O G Berg. A model for the statistical fluctuations of protein numbers in a microbial population. Journal of Theoretical Biology, 71(4):587–603, 1978.

37. D. R. Rigney and W. C. Schieve. Stochastic model of linear, continuous protein synthesis in bacterial populations. Journal of Theoretical Biology, 69(4):761–766, 1977.

38. Pavol Bokes, John R King, Andrew TA Wood, and Matthew Loose. Exact and approximate distributions of protein and mRNA levels in the low-copy regime of gene expression. Journal of Mathematical Biology, 64(5):829–854, 2012.

39. Jiajun Zhang and Tianshou Zhou. Markovian approaches to modeling intracellular reaction processes with molecular memory. Proceedings of the National Academy of Sciences, 116(47):23542–23550, 2019.

40. Zihao Wang, Zhenquan Zhang, and Tianshou Zhou. Exact distributions for stochastic models of gene expression with arbitrary regulation. arXiv preprint 1912.04680, 2019.

41. Tuoqi Liu, Jiajun Zhang, and Tianshou Zhou. Effect of interaction between chromatin loops on cell-to-cell variability in gene expression. PLoS computational biology, 12(5):e1004917, 2016.

42. Tianshou Zhou and Jiajun Zhang. Analytical results for a multistate gene model. SIAM Journal on Applied Mathematics, 72(3):789–818, 2012.

43. Nir Friedman, Long Cai, and X Sunney Xie. Linking stochastic dynamics to population distribution: an analytical framework of gene expression. Physical review letters, 97(16):168302, 2006.

44. Daphne Ezer, Victoria Moignard, Berthold Göttgens, and Boris Adryan. Determining physical mechanisms of gene expression regulation from single cell gene expression data. PLoS computational biology, 12(8):e1005072, 2016.

45. Anton JM Larsson, Per Johnsson, Michael Hagemann-Jensen, Leonard Hartmanis, Omid R Faridani, Björn Reinius, Åsa Segerstolpe, Chloe M Rivera, Bing Ren, and Rickard Sandberg. Genomic encoding of transcriptional burst kinetics. Nature, 565(7738):251, 2019.

46. Jong Kyoung Kim and John C Marioni. Inferring the kinetics of stochastic gene expression from single-cell RNA-sequencing data. Genome biology, 14(1):R7, 2013.

47. Long Cai, Nir Friedman, and X Sunney Xie. Stochastic protein expression in individual cells at the single molecule level. Nature, 440(7082):358, 2006.

48. Keren Bahar Halpern, Sivan Tanami, Shanie Landen, Michal Chapal, Liran Szlak, Anat Hutzler, Anna Nizhberg, and Shalev Itzkovitz. Bursty gene expression in the intact mammalian liver. Molecular cell, 58(1):147–156, 2015.

49. Paul J Choi, Long Cai, Kirsten Frieda, and X Sunney Xie. A stochastic single-molecule event triggers phenotype switching of a bacterial cell. Science (New York, N.Y.), 322(5900):442–6, 2008.

50. Jiajun Zhang, Qing Nie, and Tianshou Zhou. Revealing dynamic mechanisms of cell fate decisions from single-cell transcriptomic data. Frontiers in Genetics, 10(1280), 2019.

51. William Feller. An introduction to probability theory and its applications. 1957.

52. Guilherme da Costa Pereira Innocentini, Michael Forger, Alexandre Ferreira Ramos, Ovidiu Radulescu, and José Eduardo Martinho Hornos. Multimodality and flexibility of stochastic gene expression. Bulletin of mathematical biology, 75(12):2600–2630, 2013.

53. Adrienne W Kemp and CD Kemp. A family of discrete distributions defined via their factorial moments. Communications in Statistics-Theory and Methods, 3(12):1187–1196, 1974.

54. John Gurland. A generalized class of contagious distributions. Biometrics, 14(2):229–249, 1958.

55. Ram C Tripathi and John Gurland. Some aspects of the Kemp families of distributions. Communications in statistics-theory and methods, 8(9):855–869, 1979.

56. Norman Lloyd Johnson and Samuel Kotz. Urn models and their application; an approach to modern discrete probability theory. New York, NY (USA) Wiley, 1977.

57. George Pólya. Mathematics and plausible reasoning: Patterns of Plausible Inference, volume 2. Princeton University Press, 1954.

58. Hans Freudenthal. Models in applied probability. In The concept and the role of the model in mathematics and natural and social sciences, pages 78–88. Springer, 1961.

59. Lucy Ham, David Schnoerr, Rowan D Brackston, and Michael PH Stumpf. Exactly solvable models of stochastic gene expression. bioRxiv, 2020.

60. Donald A McQuarrie. Stochastic approach to chemical kinetics. Journal of applied probability, 4(3):413–478, 1967.

61. Arjun Raj, Charles S. Peskin, Daniel Tranchina, Diana Y. Vargas, and Sanjay Tyagi. Stochastic mRNA synthesis in mammalian cells. PLoS Biology, 4(10):e309, 2006.

62. Norman L Johnson, Adrienne W Kemp, and Samuel Kotz. Univariate discrete distributions, volume 444. John Wiley & Sons, 2005.

63. Johan Elf and Irmeli Barkefors. Single-molecule kinetics in living cells. Annual review of biochemistry, 88:635–659, 2019.

64. Serena Liu and Cole Trapnell. Single-cell transcriptome sequencing: recent advances and remaining challenges. F1000Research, 5, 2016.

65. Stanley Wasserman. Distinguishing between stochastic models of heterogeneity and contagion. Journal of Mathematical Psychology, 27(2):201–215, 1983.

66. John W Pearson, Sheehan Olver, and Mason A Porter. Numerical methods for the computation of the confluent and Gauss hypergeometric functions. Numerical Algorithms, 74(3):821–866, 2017.

