## Supplementary Text for "Urn models for regulated gene expression yield physically intuitive solutions for probability distributions of single-cell counts"

### 1 Distributions from models of constitutive gene expression

#### 1.1 Poisson distribution for mRNA counts

##### 1.1.1 Kinetic model

Let us consider a typical mRNA species. An elementary model of constitutive expression allows polymerase arrivals and the consequent transcription at all times [1]. An mRNA molecule is produced when an RNA polymerase arrives at the promoter for the corresponding gene. Say, the propensity for polymerase arrival is $v_0$. Let $d_0$ be the propensity of an mRNA molecule to degrade. We seek the probability, $P(E_{m_1})$ that starting from any time point when the system is at stationary state, $m_1$ polymerases arrive at the promoter in the mean lifetime of mRNA molecules, $d_0^{-1}$.

##### 1.1.2 Mapping to an urn scheme

The polymerase arrivals can be modeled as a process of unbiased sampling of balls from a well-mixed urn an infinite number of times — drawing one ball from the urn at a time, replacing it, mixing the urn, and repeating by drawing another ball [2]. The urn contains balls of black and white colors, such that the number of black balls, $n_{\text{black}}$ is infinitesimal compared to that of white balls, $n_{\text{white}}$. The black and white balls represent arrivals of RNA polymerases or other solutes, respectively.

##### 1.1.3 Relationship between parameters of the kinetic and urn models

An event of drawing a black ball from the urn corresponds to an event of arrival of an RNA polymerase. As such, we define $n_{\text{black}}$ as proportional to $v_0$. The samples are, say, drawn $n_{\text{trials}}$ times. $n_{\text{trials}}$ is infinite if we assume that the time duration for each trial, $\Delta t \to 0$. Hence, the expected value of $m_1$ is

$$\begin{aligned}\mu &= \lim_{\substack{n_{\text{trials}}\to\infty \\ n_{\text{white}}\to\infty}} \frac{n_{\text{black}}}{n_{\text{black}}+n_{\text{white}}} \cdot n_{\text{trials}} \\ &= \lim_{\Delta t\to 0} v_0 \Delta t \cdot \frac{d_0^{-1}}{\Delta t} \\ &= \frac{v_0}{d_0}. \end{aligned} \tag{1}$$

##### 1.1.4 Solution using the urn model

The sampling of balls from an urn as described above is essentially the same as a series of Bernoulli trials, such that each draw of a ball is a trial. For any trial, there are two possible outcomes — a black ball with the probability ${}^{n_{\text{black}}}/_{n_{\text{black}}+n_{\text{white}}}$ or, a white one with the probability ${}^{n_{\text{white}}}/_{n_{\text{black}}+n_{\text{white}}}$. These do not change from one trial to the next because the ball drawn in every iteration is replaced and the urn is mixed again before the next trial. The event $E_{m_1}$ of $m_1$ polymerase arrivals corresponds to the event $E_{\text{urn},m_1}$ of drawing $m_1$ black balls from the urns. Hence,

$$\begin{aligned}P(E_{m_1}) &= P(E_{\text{urn},m_1}) \\ &= \lim_{\substack{n_{\text{trials}}\to\infty \\ n_{\text{white}}\to\infty}} \binom{n_{\text{trials}}}{m_1} \left(\frac{n_{\text{black}}}{n_{\text{black}}+n_{\text{white}}}\right)^{m_1} \left(\frac{n_{\text{white}}}{n_{\text{black}}+n_{\text{white}}}\right)^{n_{\text{trials}}-m_1} \\ &= \lim_{\substack{n_{\text{trials}}\to\infty \\ n_{\text{white}}\to\infty}} \frac{n_{\text{trials}}!}{m_1!\,(n_{\text{trials}}-m_1)!} \left(\frac{n_{\text{black}}}{n_{\text{black}}+n_{\text{white}}}\right)^{m_1} \left(\frac{n_{\text{white}}}{n_{\text{black}}+n_{\text{white}}}\right)^{n_{\text{trials}}-m_1} \\ &= \frac{1}{m_1!} \lim_{\substack{n_{\text{trials}}\to\infty \\ n_{\text{white}}\to\infty}} \left(\frac{n_{\text{white}}}{n_{\text{black}}+n_{\text{white}}}\right)^{-m_1} \left(1-\frac{n_{\text{black}}}{n_{\text{black}}+n_{\text{white}}}\right)^{n_{\text{trials}}} \\ &\qquad \left\{\frac{n_{\text{trials}}(n_{\text{trials}}-1)(n_{\text{trials}}-2)\ldots(n_{\text{trials}}-m_1+1)}{(n_{\text{black}}+n_{\text{white}})^{m_1}} n_{\text{black}}^{m_1}\right\} \\ &= \frac{1}{m_1!} \lim_{\substack{n_{\text{trials}}\to\infty \\ n_{\text{white}}\to\infty}} \left(1-\frac{n_{\text{black}}}{n_{\text{black}}+n_{\text{white}}}\right)^{n_{\text{trials}}} \left(\frac{n_{\text{trials}} n_{\text{black}}}{n_{\text{black}}+n_{\text{white}}}\right)^{m_1} \\ &= \frac{1}{m_1!} \lim_{\Delta t\to 0} (1-v_0\Delta t)^{d_0^{-1}/\Delta t} \left(v_0\Delta t \cdot \frac{d_0^{-1}}{\Delta t}\right)^{m_1} \\ &\qquad \left[\text{using } e = \lim_{x\to 0}(1+x)^{\frac{1}{x}}\right] \\ &= \frac{e^{-\mu}\mu^{m_1}}{m_1!}. \end{aligned}$$

We denote this distribution $\text{Pois}(\bullet_{m_1}|\mu)$.

#### 1.2 Negative binomial distribution for protein counts when mRNAs decay much faster than proteins

##### 1.2.1 Kinetic model

One of the elementary models for constitutive protein production builds on the kinetic model described in the previous section. In this model, the proteins are produced by translation of mRNA and ribosomes function as catalysts. Ribosomes arrive on an mRNA molecule with a uniform propensity of $v_1$ and a protein molecule decays with the propensity $d_1$ [1, 3]. $v_0$ and $d_0$ are defined as in the previous section. We seek the probability, $P(E_{m_2})$ that starting from a time when the system is at stationary state, $m_2$ ribosomes arrive on mRNAs produced in $d_1^{-1}$ time. Typically, the mRNAs decay much faster than the proteins, i.e., $d_0 \gg d_1$ [3]. Then, the kinetic scheme is approximated as a process of bursty production of proteins, whereby at exponentially distributed time intervals, several proteins are produced simultaneously.

###### 1.2.2 Mapping to an urn scheme

The process of translation can be modeled as one of the outcomes in a sampling of balls with replacement from a well-mixed urn with finite numbers of balls of black and white colors. An outcome of black ball means that a ribosome arrives on an mRNA and translation occurs. An outcome of white ball represents the arrival of an RNAse, which degrades the mRNA molecule. Once, again, let $n_{\text{black}}$ and $n_{\text{white}}$ be the numbers of the black and the white balls in the urn. Unlike the urn scheme for transcription, for translation, the process of sampling stops after first outcome of white ball, which represents the arrival of an RNAse leading to decay of the mRNA.

###### 1.2.3 Relationship between parameters of the kinetic and urn models

Drawing a black ball corresponds to a ribosome arrival. Hence, $n_{\text{black}}$ is proportional to $v_1$. Starting from any time point, the next event in an mRNA molecule's life could either be its decay or its translation by ribosome. The probability of a ribosome arrival is $n_{\text{black}}/n_{\text{black}}+n_{\text{white}} = v_1/v_1+d_0$. Let $\beta = v_1/d_0$ be the expected number of ribosome arrivals per mRNA molecule, also called the burst size. Then, $n_{\text{black}}/n_{\text{black}}+n_{\text{white}} = \beta/1+\beta$.

Given that $d_0 \gg d_1$, the noise in mRNA distribution exists on too fast a time scale to be observed over protein lifetimes. Hence, we work with the mean number of RNA polymerase arrivals in a protein lifetime, i.e., burst frequency as sufficient statistic for the contribution of transcription to noise in translation. Let this number be $\alpha = v_0/d_1$.

###### 1.2.4 Solution using the urn model

Once again, we sample balls from the urn in a series of Bernoulli trials. We stop when the first white ball appears (mRNA decay) and record the number of black balls until the first white ball as the outcome of interest. This is the number of ribosome arrivals per mRNA and hence, of the proteins produced.

For each polymerase arrival, we sample balls from the urn as described above. Let $m_2$ be the cumulative count of ribosome arrivals for all the polymerase arrivals in a protein lifetime. An event, $E_{m_2}$ of $m_2$ ribosome arrivals corresponds to an event, $E_{\text{urn},m_2}$ of $m_2$ black balls separated by $\alpha$ white balls, such that the last ball is always white. Hence,

$$\begin{aligned} P(E_{m_2}) &= P(E_{\text{urn},m_2}) \\ &= \binom{\alpha+m_2-1}{m_2}\left(\frac{n_{\text{black}}}{n_{\text{black}}+n_{\text{white}}}\right)^{m_2}\left(1-\frac{n_{\text{black}}}{n_{\text{black}}+n_{\text{white}}}\right)^{\alpha} \\ &= \binom{\alpha+m_2-1}{m_2}\left(\frac{\beta}{1+\beta}\right)^{m_2}\left(1-\frac{\beta}{1+\beta}\right)^{\alpha}. \end{aligned}$$

We denote this distribution $\text{NB}(\bullet_{m_2}|\alpha,\beta)$.

#### 2 Negative hypergeometric distribution

Negative hypergeometric distribution can result from a variety of urn schemes [4]. One of the popular ways to derive it is by considering a scheme where balls are drawn from an urn but not replaced [5, 6]. For use in the main text, we derive it using a scheme of assigning a given number of identical balls to a set of urns of two different colors. Below, we describe both the schemes.

##### 2.1 Derivation based on a scheme of sampling balls without replacement from an urn with black and white balls until $m$ black balls are drawn

###### 2.1.1 The urn scheme

Let us consider an urn that contains a finite number of black and white balls. Once again, we use the notation $n_{\text{black}}$ and $n_{\text{white}}$ for the black and white balls. If we draw balls from the urn without replacing them, the probability of drawing either of the two colors varies with the composition of the urn from one draw to the next. Let the outcome of interest be the number of white balls, $i$ drawn until the $m^{\text{th}}$ black ball is obtained, with $m < n_{\text{black}}$.

###### 2.1.2 Solution

Of the $m+i$ draws, the first $m+i-1$ must contain $m-1$ black balls in any order, and the last draw must be a black ball. Hence, the probability, $p(i)$ of an outcome $i$ is

$$\begin{aligned} p(i) &= \frac{\binom{n_{\text{white}}}{i}\binom{n_{\text{black}}}{m-1}}{\binom{n_{\text{black}}+n_{\text{white}}}{m+i-1}}\frac{n_{\text{black}}-m+1}{n_{\text{black}}+n_{\text{white}}-m-i+1} \\ &= \frac{\binom{m+i-1}{m-1}\binom{n_{\text{black}}+n_{\text{white}}-m-i}{n_{\text{black}}-m}}{\binom{n_{\text{black}}+n_{\text{white}}}{n_{\text{black}}}}. \end{aligned} \tag{2}$$

##### 2.2 Derivation based on a scheme of assigning a given number of balls to a set of red and grey urns such that $m$ of them are assigned to the red urns

###### 2.2.1 The urn scheme

Our urn scheme involves assigning balls to a set of urns instead of drawing from an urn. In our scheme, the urns are colored — red or grey, and the balls are all identical, say black in color. Let us say that we have $n_{\text{red}}$ red urns and $n_{\text{grey}}$ grey urns. For our urn experiment, we shall assign a given number, $m+i$ of black balls to these urns. Our outcome of interest is the number of balls, $m$ that are assigned to the red urns. Additional features of our scheme are worth defining — the urns are identical in sizes, arranged in a row and distinguishable by their positions, can contain an infinite number of balls, empty urns are allowed and all balls are assigned randomly to the urns irrespective of their colors or positions.

###### 2.2.2 Solution

First, note that the number of ways to distribute $m$ balls in say, $n_{\text{urns}}$, all of the same kind is given by $\binom{n_{\text{urns}}+m-1}{m}$ (see Table 1.2, page 37 of Ref. [2]). To see this, consider the balls along with $m-1$ dividers. Every ordered arrangement of the balls and the dividers is one way of dividing the balls in the urns, with the balls between two dividers going to a separate urn. More specifically, the balls between the $k^{\text{th}}$ and $(k+1)^{\text{th}}$ dividers go to the $(k+1)^{\text{th}}$ urn. Also, the balls to the left of the first divider and to the right of the last divider go to the first and the last urns. Since there are $\binom{n_{\text{urns}}+m-1}{m}$ ways of arranging the balls and the dividers, there are the same number of ways to assign $m$ balls to $n_{\text{urns}}$ urns.

Given this result, we can derive the probability of assigning $m+i$ balls to $n_{\text{red}}+n_{\text{grey}}$ urns such that $m$ balls are assigned to the red urns. In fact, this is simply the ratio of the following two quantities — (a) the number of ways to assign $m$ and $i$ balls to the red and grey urns, respectively and, (b) the number of ways to assign $m+i$ balls to any of the urns, i.e.,

$$p(m) = \frac{\binom{n_{\text{red}}+m-1}{m}\binom{n_{\text{grey}}+i-1}{i}}{\binom{n_{\text{red}}+n_{\text{grey}}+m+i-1}{m+i}}. \tag{3}$$

We denote this distribution $\text{NH}\left(\bullet_m \mapsto n_{\text{red}} | \bullet_{m+i} \mapsto \{n_{\text{red}}, n_{\text{grey}}\}\right)$. Eqs. 2 and 3 are formally identical. In fact, by substituting the terms in Eq. 3 as $m \equiv i$, $i \equiv n_{\text{white}} - i$, $n_{\text{grey}} \equiv n_{\text{black}} - m + 1$ and $n_{\text{red}} \equiv m$, we can retrieve Eq. 2.

#### 3 Proofs showing that the solutions from the urn model approach and chemical master equations are identical

In the following, $\Gamma(x)$ represents the gamma function and equals $(x-1)!$ when $x$ is a positive integer. Also, $(a)_i$ is the Pocchammer symbol for the ascending factorial. It is defined as $(a)_i = \frac{\Gamma(a+i)}{\Gamma(a)}$.

##### 3.1 Peccoud-Ycart model [7]

$$
\begin{aligned}
P\left(E_{\text{urn},m_1}\right) &= P\left(\bullet_{m_1} \mapsto n_{\text{red}} | \mu, n_{\text{red}}, n_{\text{grey}}\right) \\
&= \sum_{i_1=0}^{\infty} \text{Pois}\left(\bullet_{m_1+i_1} | \mu\right) \times \text{NH}\left(\bullet_{m_1} \mapsto n_{\text{red}} | \bullet_{m_1+i_1} \mapsto \{n_{\text{red}}, n_{\text{grey}}\}\right) \\
&= e^{-\mu} \sum_{i_1=0}^{\infty} \frac{\binom{n_{\text{red}}+m_1-1}{m_1}\binom{n_{\text{grey}}+i_1-1}{i_1}}{\binom{n_{\text{red}}+n_{\text{grey}}+m_1+i_1-1}{m_1+i_1}} \frac{\mu^{m_1+i_1}}{(m_1+i_1)!} \\
&= e^{-\mu} \sum_{i_1=0}^{\infty} \frac{\frac{(n_{\text{red}}+m_1-1)!}{(n_{\text{red}}-1)!m_1!}\frac{(n_{\text{grey}}+i_1-1)!}{(n_{\text{grey}}-1)!i_1!}}{\frac{(n_{\text{red}}+n_{\text{grey}}+m_1+i_1-1)!}{(n_{\text{red}}+n_{\text{grey}}-1)!(m_1+i_1)!}} \frac{\mu^{m_1+i_1}}{(m_1+i_1)!} \\
&= \frac{e^{-\mu}\mu^{m_1}}{m_1!} \frac{\Gamma\left(n_{\text{red}}+m_1\right)}{\Gamma\left(n_{\text{red}}\right)} \frac{\Gamma\left(n_{\text{red}}+n_{\text{grey}}\right)}{\Gamma\left(n_{\text{red}}+n_{\text{grey}}+m_1\right)} \sum_{i_1=0}^{\infty} \frac{\Gamma\left(n_{\text{grey}}+i_1\right)}{\Gamma\left(n_{\text{grey}}\right)} \frac{\Gamma\left(n_{\text{red}}+n_{\text{grey}}+m_1\right)}{\Gamma\left(n_{\text{red}}+n_{\text{grey}}+m_1+i_1\right)} \frac{\mu^{i_1}}{i_1!} \\
&= \frac{\left(n_{\text{red}}\right)_{m_1}}{\left(n_{\text{red}}+n_{\text{grey}}\right)_{m_1}} \frac{\mu^{m_1}}{m_1!} e^{-\mu} \sum_{i_1=0}^{\infty} \frac{\left(n_{\text{grey}}\right)_{i_1}}{\left(n_{\text{red}}+n_{\text{grey}}+m_1\right)_{i_1}} \frac{\mu^{i_1}}{i_1!} \\
&= \frac{\left(n_{\text{red}}\right)_{m_1}}{\left(n_{\text{red}}+n_{\text{grey}}\right)_{m_1}} \frac{\mu^{m_1}}{m_1!} e^{-\mu} {}_1F_1\left[n_{\text{grey}}; n_{\text{red}}+n_{\text{grey}}+m_1; \mu\right] \\
&\qquad\qquad \text{[Using Kummer transformation in Eq. 13.1.27 of Ref. [8]]} \\
&= \frac{\left(n_{\text{red}}\right)_{m_1}}{\left(n_{\text{red}}+n_{\text{grey}}\right)_{m_1}} \frac{\mu^{m_1}}{m_1!} {}_1F_1\left[n_{\text{red}}+m_1; n_{\text{red}}+n_{\text{grey}}+m_1; -\mu\right] \\
&= \frac{1}{m_1!} \frac{d^{m_1}}{dz^{m_1}} {}_1F_1\left[n_{\text{red}}; n_{\text{red}}+n_{\text{grey}}; \mu(z-1)\right]\Bigg|_{z=0} \\
&= \text{Coefficient of } z^{m_1} \text{ in the Maclaurin series expansion of } {}_1F_1\left[n_{\text{red}}; n_{\text{red}}+n_{\text{grey}}; \mu(z-1)\right] \\
&= P\left(E_{m_1}\right)
\end{aligned}
$$

##### 3.2 Shahrezaei-Swain model [3]

$$\begin{aligned}
P\left(E_{\mathrm{urn},m_2}\right) &= P\left(\bullet_{m_2} \mapsto n_{\mathrm{red}} | \alpha, \beta, n_{\mathrm{red}}, n_{\mathrm{grey}}\right) \\
&= \sum_{i_2=0}^{\infty} \mathrm{NB}\left(\bullet_{m_2+i_2} | \alpha, \beta\right) \times \mathrm{NH}\left(\bullet_{m_2} \mapsto n_{\mathrm{red}} | \bullet_{m_2+i_2} \mapsto \{n_{\mathrm{red}}, n_{\mathrm{grey}}\}\right) \\
&= \sum_{i_2=0}^{\infty} \frac{\binom{n_{\mathrm{red}}+m_2-1}{m_2}\binom{n_{\mathrm{grey}}+i_2-1}{i_2}}{\binom{n_{\mathrm{red}}+n_{\mathrm{grey}}+m_2+i_2-1}{m_2+i_2}} \left[\binom{\alpha+m_2+i_2-1}{m_2+i_2}\left(\frac{1}{1+\beta}\right)^{\alpha}\left(\frac{\beta}{1+\beta}\right)^{m_2+i_2}\right] \\
&= \sum_{i_2=0}^{\infty} \frac{\frac{(n_{\mathrm{red}}+m_2-1)!}{(n_{\mathrm{red}}-1)!m_2!}\frac{(n_{\mathrm{grey}}+i_2-1)!}{(n_{\mathrm{grey}}-1)!i_2!}}{\frac{(n_{\mathrm{red}}+n_{\mathrm{grey}}+m_2+i_2-1)!}{(n_{\mathrm{red}}+n_{\mathrm{grey}}-1)!(m_2+i_2)!}} \left[\frac{(\alpha+m_2+i_2-1)!}{(\alpha-1)!\,(m_2+i_2)!}\left(\frac{1}{1+\beta}\right)^{\alpha}\left(\frac{\beta}{1+\beta}\right)^{m_2+i_2}\right] \\
&= \frac{1}{m_2!}\frac{\Gamma(\alpha+m_2)}{\Gamma(\alpha)}\frac{\Gamma(n_{\mathrm{red}}+m_2)}{\Gamma(n_{\mathrm{red}})}\frac{\Gamma(n_{\mathrm{red}}+n_{\mathrm{grey}})}{\Gamma(n_{\mathrm{red}}+n_{\mathrm{grey}}+m_2)}\left(\frac{1}{1+\beta}\right)^{\alpha}\left(\frac{\beta}{1+\beta}\right)^{m_2} \\
&\quad \times \sum_{i_2=0}^{\infty}\frac{\Gamma(\alpha+m_2+i_2)}{\Gamma(\alpha+m_2)}\frac{\Gamma(n_{\mathrm{grey}}+i_2)}{\Gamma(n_{\mathrm{grey}})}\frac{\Gamma(n_{\mathrm{red}}+n_{\mathrm{grey}}+m_2)}{\Gamma(n_{\mathrm{red}}+n_{\mathrm{grey}}+m_2+i_2)}\left(\frac{\beta}{1+\beta}\right)^{i_2}\frac{1}{i_2!} \\
&= \frac{1}{m_2!}\frac{(\alpha)_{m_2}(n_{\mathrm{red}})_{m_2}}{(n_{\mathrm{red}}+n_{\mathrm{grey}})_{m_2}}\left(\frac{1}{1+\beta}\right)^{\alpha}\left(\frac{\beta}{1+\beta}\right)^{m_2}\sum_{i_2=0}^{\infty}\frac{(\alpha+m_2)_{i_2}(n_{\mathrm{grey}})_{i_2}}{(n_{\mathrm{red}}+n_{\mathrm{grey}}+m_2)_{i_2}}\left(\frac{\beta}{1+\beta}\right)^{i_2}\frac{1}{i_2!} \\
&= \frac{1}{m_2!}\frac{(\alpha)_{m_2}(n_{\mathrm{red}})_{m_2}}{(n_{\mathrm{red}}+n_{\mathrm{grey}})_{m_2}}\left(\frac{1}{1+\beta}\right)^{\alpha}\left(\frac{\beta}{1+\beta}\right)^{m_2}{}_2F_1\left[\alpha+m_2, n_{\mathrm{grey}}; n_{\mathrm{red}}+n_{\mathrm{grey}}+m_2; \frac{\beta}{1+\beta}\right] \\
&\qquad\qquad\qquad\qquad\qquad\qquad \text{[Using the transformation formula in Eq. 15.3.4 of Ref. [8]]} \\
&= \frac{(\alpha)_{m_2}(n_{\mathrm{red}})_{m_2}}{(n_{\mathrm{red}}+n_{\mathrm{grey}})_{m_2}}\frac{\beta^{m_2}}{m_2!}\;{}_2F_1\left[\alpha+m_2, n_{\mathrm{red}}+m_2; n_{\mathrm{red}}+n_{\mathrm{grey}}+m_2; -\beta\right] \\
&= \frac{1}{m_2!}\frac{d^{m_2}}{dz^{m_2}}{}_2F_1\left[\alpha, n_{\mathrm{red}}; n_{\mathrm{red}}+n_{\mathrm{grey}}; \beta(z-1)\right]\Big|_{z=0} \\
&= \text{Coefficient of } z^{m_2} \text{ in the Maclaurin series expansion of } {}_2F_1\left[\alpha, n_{\mathrm{red}}; n_{\mathrm{red}}+n_{\mathrm{grey}}; \beta(z-1)\right] \\
&= P\left(E_{m_2}\right)
\end{aligned}$$

##### 3.3 The three-state model for mRNAs by Cao *et al.* [9] and Karmakar [10]

$$\begin{aligned}
P\left(E_{\mathrm{urn},m_1}\right) &= P\left(\bullet_{m_1} \mapsto n_{\mathrm{red}} | \mu, n_{\mathrm{red}}, n_{\mathrm{grey}}, n_{\mathrm{ppl}}, n_{\mathrm{blue}}\right) \\
&= \sum_{i_1=0}^{\infty}\sum_{j_1=0}^{\infty} \mathrm{Pois}\left(\bullet_{m_1+i_1+j_1}|\mu\right) \times \mathrm{NH}\left(\bullet_{m_1+i_1} \mapsto n_{\mathrm{ppl}} | \bullet_{m_1+i_1+j_1} \mapsto \{n_{\mathrm{ppl}}, n_{\mathrm{blue}}\}\right) \times \\
&\qquad \mathrm{NH}\left(\bullet_{m_1} \mapsto n_{\mathrm{red}} | \bullet_{m_1+i_1} \mapsto \{n_{\mathrm{red}}, n_{\mathrm{grey}}\}\right) \\
&= \sum_{i_1=0}^{\infty}\sum_{j_1=0}^{\infty} \frac{\binom{n_{\mathrm{red}}+m_1-1}{m_1}\binom{n_{\mathrm{grey}}+i_1-1}{i_1}}{\binom{n_{\mathrm{red}}+n_{\mathrm{grey}}+m_1+i_1-1}{m_1+i_1}} \cdot \frac{\binom{n_{\mathrm{ppl}}+m_1+i_1-1}{m_1+i_1}\binom{n_{\mathrm{blue}}+j_1-1}{j_1}}{\binom{n_{\mathrm{ppl}}+n_{\mathrm{blue}}+m_1+i_1+j_1-1}{m_1+i_1+j_1}} \cdot \frac{e^{-\mu}\mu^{m_1+i_1+j_1}}{(m_1+i_1+j_1)!} \\
&= \sum_{i_1=0}^{\infty}\sum_{j_1=0}^{\infty} \frac{\frac{(n_{\mathrm{red}}+m_1-1)!}{(n_{\mathrm{red}}-1)!m_1!}\frac{(n_{\mathrm{grey}}+i_1-1)!}{(n_{\mathrm{grey}}-1)!i_1!}}{\frac{(n_{\mathrm{red}}+n_{\mathrm{grey}}+m_1+i_1-1)!}{(n_{\mathrm{red}}+n_{\mathrm{grey}}-1)!(m_1+i_1)!}} \cdot \frac{\frac{(n_{\mathrm{ppl}}+m_1+i_1-1)!}{(n_{\mathrm{ppl}}-1)!(m_1+i_1)!}\frac{(n_{\mathrm{blue}}+j_1-1)!}{(n_{\mathrm{blue}}-1)!j_1!}}{\frac{(n_{\mathrm{ppl}}+n_{\mathrm{blue}}+m_1+i_1+j_1-1)!}{(n_{\mathrm{ppl}}+n_{\mathrm{blue}}-1)!(m_1+i_1+j_1)!}} \cdot \frac{e^{-\mu}\mu^{m_1+i_1+j_1}}{(m_1+i_1+j_1)!} \\
&= \sum_{i_1=0}^{\infty}\sum_{j_1=0}^{\infty} \frac{\left(n_{\mathrm{red}}\right)_{m_1}\left(n_{\mathrm{grey}}\right)_{i_1}}{\left(n_{\mathrm{red}}+n_{\mathrm{grey}}\right)_{m_1+i_1}} \cdot \frac{\left(n_{\mathrm{ppl}}\right)_{m_1+i_1}\left(n_{\mathrm{blue}}\right)_{j_1}}{\left(n_{\mathrm{ppl}}+n_{\mathrm{blue}}\right)_{m_1+i_1+j_1}} \cdot \frac{e^{-\mu}\mu^{m_1+i_1+j_1}}{m_1!i_1!j_1!} \\
&\qquad\qquad \left[\text{Using } (a)_{x+y} = (a)_x (a+x)_y\right] \\
&= \sum_{i_1=0}^{\infty}\sum_{j_1=0}^{\infty} \frac{\left(n_{\mathrm{red}}\right)_{m_1}\left(n_{\mathrm{grey}}\right)_{i_1}}{\left(n_{\mathrm{red}}+n_{\mathrm{grey}}\right)_{m_1}\left(n_{\mathrm{red}}+n_{\mathrm{grey}}+m_1\right)_{i_1}} \cdot \\
&\qquad \frac{\left(n_{\mathrm{ppl}}\right)_{m_1}\left(n_{\mathrm{ppl}}+m_1\right)_{i_1}\left(n_{\mathrm{blue}}\right)_{j_1}}{\left(n_{\mathrm{ppl}}+n_{\mathrm{blue}}\right)_{m_1}\left(n_{\mathrm{ppl}}+n_{\mathrm{blue}}+m_1\right)_{j_1}\left(n_{\mathrm{ppl}}+n_{\mathrm{blue}}+m_1+j_1\right)_{i_1}} \cdot \\
&\qquad \frac{e^{-\mu}\mu^{m_1+i_1+j_1}}{m_1!i_1!j_1!} \\
&= \frac{\left(n_{\mathrm{red}}\right)_{m_1}\left(n_{\mathrm{ppl}}\right)_{m_1}}{\left(n_{\mathrm{red}}+n_{\mathrm{grey}}\right)_{m_1}\left(n_{\mathrm{ppl}}+n_{\mathrm{blue}}\right)_{m_1}} \cdot \frac{e^{-\mu}\mu^{m_1}}{m_1!} \cdot \\
&\qquad \sum_{j_1=0}^{\infty} \frac{\left(n_{\mathrm{blue}}\right)_{j_1}}{\left(n_{\mathrm{ppl}}+n_{\mathrm{blue}}+m_1\right)_{j_1}} \cdot \frac{\mu^{j_1}}{j_1!} \cdot \\
&\qquad\qquad \sum_{i_1=0}^{\infty} \frac{\left(n_{\mathrm{grey}}\right)_{i_1}\left(n_{\mathrm{ppl}}+m_1\right)_{i_1}}{\left(n_{\mathrm{red}}+n_{\mathrm{grey}}+m_1\right)_{i_1}\left(n_{\mathrm{ppl}}+n_{\mathrm{blue}}+m_1+j_1\right)_{i_1}} \frac{\mu^{i_1}}{i_1!} \\
&= \frac{\left(n_{\mathrm{red}}\right)_{m_1}\left(n_{\mathrm{ppl}}\right)_{m_1}}{\left(n_{\mathrm{red}}+n_{\mathrm{grey}}\right)_{m_1}\left(n_{\mathrm{ppl}}+n_{\mathrm{blue}}\right)_{m_1}} \cdot \frac{\mu^{m_1}}{m_1!} \cdot \\
&\qquad e^{-\mu}\sum_{j_1=0}^{\infty} \frac{\left(n_{\mathrm{blue}}\right)_{j_1}}{\left(n_{\mathrm{ppl}}+n_{\mathrm{blue}}+m_1\right)_{j_1}} \cdot \frac{\mu^{j_1}}{j_1!} \cdot \\
&\qquad\qquad {}_2F_2\left[{n_{\mathrm{grey}},\ n_{\mathrm{ppl}}+m_1 \atop n_{\mathrm{red}}+n_{\mathrm{grey}}+m_1,\ n_{\mathrm{ppl}}+n_{\mathrm{blue}}+m_1+j_1}; \mu\right] \\
&\qquad\qquad [\text{Using the Kummer-type transformation in Eq. 3 of Paris, 2005 [11]}] \\
&= \frac{\left(n_{\mathrm{red}}\right)_{m_1}\left(n_{\mathrm{ppl}}\right)_{m_1}}{\left(n_{\mathrm{red}}+n_{\mathrm{grey}}\right)_{m_1}\left(n_{\mathrm{ppl}}+n_{\mathrm{blue}}\right)_{m_1}} \cdot \frac{\mu^{m_1}}{m_1!} \cdot \\
&\qquad {}_2F_2\left[{n_{\mathrm{red}}+m_1,\ n_{\mathrm{ppl}}+m_1 \atop n_{\mathrm{red}}+n_{\mathrm{grey}}+m_1,\ n_{\mathrm{ppl}}+n_{\mathrm{blue}}+m_1}; -\mu\right] \\
&= \frac{1}{m_1!}\frac{d^{m_1}}{dz^{m_1}}\, {}_2F_2\left[{n_{\mathrm{red}},\ n_{\mathrm{ppl}} \atop n_{\mathrm{red}}+n_{\mathrm{grey}},\ n_{\mathrm{ppl}}+n_{\mathrm{blue}}}; \mu(z-1)\right]\Bigg|_{z=0} \\
&= \text{Coefficient of } z^{m_1} \text{ in the Maclaurin series expansion of } {}_2F_2\left[{n_{\mathrm{red}},\ n_{\mathrm{ppl}} \atop n_{\mathrm{red}}+n_{\mathrm{grey}},\ n_{\mathrm{ppl}}+n_{\mathrm{blue}}}; \mu(z-1)\right] \\
&= P\left(E_{m_1}\right)
\end{aligned}$$

##### 3.4 The three-state model for proteins by Cao *et al.* [9]

$$
\begin{aligned}
P\left(E_{\text{urn},m_2}\right) &= P\left(\bullet_{m_2} \mapsto n_{\text{red}} | \alpha, \beta, n_{\text{red}}, n_{\text{grey}}, n_{\text{ppl}}, n_{\text{blue}}\right) \\
&= \sum_{i_2=0}^{\infty}\sum_{j_2=0}^{\infty} \text{NB}\left(\bullet_{m_2+i_2+j_2}|\alpha,\beta\right) \times \text{NH}\left(\bullet_{m_2+i_2} \mapsto n_{\text{ppl}} | \bullet_{m_2+i_2+j_2} \mapsto \{n_{\text{ppl}}, n_{\text{blue}}\}\right) \times \\
&\qquad \text{NH}\left(\bullet_{m_2} \mapsto n_{\text{red}} | \bullet_{m_2+i_2} \mapsto \{n_{\text{red}}, n_{\text{grey}}\}\right) \\
&= \sum_{i_2=0}^{\infty}\sum_{j_2=0}^{\infty} \frac{\binom{n_{\text{red}}+m_2-1}{m_2}\binom{n_{\text{grey}}+i_2-1}{i_2}}{\binom{n_{\text{red}}+n_{\text{grey}}+m_2+i_2-1}{m_2+i_2}} \cdot \frac{\binom{n_{\text{ppl}}+m_2+i_2-1}{m_2+i_2}\binom{n_{\text{blue}}+j_2-1}{j_2}}{\binom{n_{\text{ppl}}+n_{\text{blue}}+m_2+i_2+j_2-1}{m_2+i_2+j_2}} \cdot \\
&\qquad \left[\frac{(\alpha+m_2+i_2+j_2-1)!}{(\alpha-1)!\,(m_2+i_2+j_2)!}\left(\frac{1}{1+\beta}\right)^{\alpha}\left(\frac{\beta}{1+\beta}\right)^{m_2+i_2+j_2}\right] \\
&= \sum_{i_2=0}^{\infty}\sum_{j_2=0}^{\infty} \frac{\frac{(n_{\text{red}}+m_2-1)!}{(n_{\text{red}}-1)!m_2!}\frac{(n_{\text{grey}}+i_2-1)!}{(n_{\text{grey}}-1)!i_2!}}{\frac{(n_{\text{red}}+n_{\text{grey}}+m_2+i_2-1)!}{(n_{\text{red}}+n_{\text{grey}}-1)!(m_2+i_2)!}} \cdot \frac{\frac{(n_{\text{ppl}}+m_2+i_2-1)!}{(n_{\text{ppl}}-1)!(m_2+i_2)!}\frac{(n_{\text{blue}}+j_2-1)!}{(n_{\text{blue}}-1)!j_2!}}{\frac{(n_{\text{ppl}}+n_{\text{blue}}+m_2+i_2+j_2-1)!}{(n_{\text{ppl}}+n_{\text{blue}}-1)!(m_2+i_2+j_2)!}} \cdot \\
&\qquad \left[\frac{(\alpha+m_2+i_2+j_2-1)!}{(\alpha-1)!\,(m_2+i_2+j_2)!}\left(\frac{1}{1+\beta}\right)^{\alpha}\left(\frac{\beta}{1+\beta}\right)^{m_2+i_2+j_2}\right] \\
&= \sum_{i_2=0}^{\infty}\sum_{j_2=0}^{\infty} \frac{(n_{\text{red}})_{m_2}(n_{\text{grey}})_{i_2}}{(n_{\text{red}}+n_{\text{grey}})_{m_2+i_2}} \cdot \frac{(n_{\text{ppl}})_{m_2+i_2}(n_{\text{blue}})_{j_2}}{(n_{\text{ppl}}+n_{\text{blue}})_{m_2+i_2+j_2}} \cdot \left[\frac{(\alpha)_{m_2+i_2+j_2}}{m_2!i_2!j_2!}\left(\frac{1}{1+\beta}\right)^{\alpha}\left(\frac{\beta}{1+\beta}\right)^{m_2+i_2+j_2}\right] \\
&\qquad \text{[Using } (a)_{x+y} = (a)_x\,(a+x)_y\text{]} \\
&= \sum_{i_2=0}^{\infty}\sum_{j_2=0}^{\infty} \frac{(n_{\text{red}})_{m_2}(n_{\text{grey}})_{i_2}}{(n_{\text{red}}+n_{\text{grey}})_{m_2}(n_{\text{red}}+n_{\text{grey}}+m_2)_{i_2}} \cdot \\
&\qquad \frac{(n_{\text{ppl}})_{m_2}(n_{\text{ppl}}+m_2)_{i_2}(n_{\text{blue}})_{j_2}}{(n_{\text{ppl}}+n_{\text{blue}})_{m_2}(n_{\text{ppl}}+n_{\text{blue}}+m_2)_{j_2}(n_{\text{ppl}}+n_{\text{blue}}+m_2+j_2)_{i_2}} \cdot \\
&\qquad \left[\frac{(\alpha)_{m_2}(\alpha+m_2)_{j_2}(\alpha+m_2+j_2)_{i_2}}{m_2!i_2!j_2!}\left(\frac{1}{1+\beta}\right)^{\alpha}\left(\frac{\beta}{1+\beta}\right)^{m_2+i_2+j_2}\right] \\
&= \frac{(\alpha)_{m_2}(n_{\text{red}})_{m_2}(n_{\text{ppl}})_{m_2}}{(n_{\text{red}}+n_{\text{grey}})_{m_2}(n_{\text{ppl}}+n_{\text{blue}})_{m_2}} \cdot \frac{\beta^{m_2}}{m_2!} \cdot \\
&\qquad \left(\frac{1}{1+\beta}\right)^{\alpha+m_2} \sum_{j_2=0}^{\infty} \frac{(\alpha+m_2)_{j_2}(n_{\text{blue}})_{j_2}}{(n_{\text{ppl}}+n_{\text{blue}}+m_2)_{j_2}} \cdot \frac{1}{j_2!}\left(\frac{\beta}{1+\beta}\right)^{j_2} \cdot \\
&\qquad \sum_{i_2=0}^{\infty} \frac{(\alpha+m_2+j_2)_{i_2}(n_{\text{grey}})_{i_2}(n_{\text{ppl}}+m_2)_{i_2}}{(n_{\text{red}}+n_{\text{grey}}+m_2)_{i_2}(n_{\text{ppl}}+n_{\text{blue}}+m_2+j_2)_{i_2}} \frac{1}{i_2!}\left(\frac{\beta}{1+\beta}\right)^{i_2} \\
&= \frac{(\alpha)_{m_2}(n_{\text{red}})_{m_2}(n_{\text{ppl}})_{m_2}}{(n_{\text{red}}+n_{\text{grey}})_{m_2}(n_{\text{ppl}}+n_{\text{blue}})_{m_2}} \cdot \frac{\beta^{m_2}}{m_2!} \cdot \\
&\qquad \left(\frac{1}{1+\beta}\right)^{\alpha+m_2} \sum_{j_2=0}^{\infty} \frac{(\alpha+m_2)_{j_2}(n_{\text{blue}})_{j_2}}{(n_{\text{ppl}}+n_{\text{blue}}+m_2)_{j_2}} \cdot \frac{1}{j_2!}\left(\frac{\beta}{1+\beta}\right)^{j_2} \cdot \\
&\qquad {}_3F_2\left[\begin{matrix}\alpha+m_2+j_2, & n_{\text{grey}}, & n_{\text{ppl}}+m_2 \\ & n_{\text{red}}+n_{\text{grey}}+m_2, & n_{\text{ppl}}+n_{\text{blue}}+m_2+j_2\end{matrix}; \frac{\beta}{1+\beta}\right] \\
&\qquad \text{[Using the Euler-type transformation in Theorem 4 of Choudhary, 2020 [12]]} \\
&= \frac{(\alpha)_{m_2}(n_{\text{red}})_{m_2}(n_{\text{ppl}})_{m_2}}{(n_{\text{red}}+n_{\text{grey}})_{m_2}(n_{\text{ppl}}+n_{\text{blue}})_{m_2}} \cdot \frac{\beta^{m_2}}{m_2!} \cdot \\
&\qquad {}_3F_2\left[\begin{matrix}\alpha+m_2, & n_{\text{red}}+m_2, & n_{\text{ppl}}+m_2 \\ & n_{\text{red}}+n_{\text{grey}}+m_2, & n_{\text{ppl}}+n_{\text{blue}}+m_2\end{matrix}; -\beta\right] \\
&= \frac{1}{m_2!}\frac{d^{m_2}}{dz^{m_2}} {}_3F_2\left[\begin{matrix}\alpha, & n_{\text{red}}, & n_{\text{ppl}} \\ & n_{\text{red}}+n_{\text{grey}}, & n_{\text{ppl}}+n_{\text{blue}}\end{matrix}; \beta(z-1)\right]\Bigg|_{z=0} \\
&= \text{Coefficient of } z^{m_2} \text{ in the Maclaurin series expansion of } {}_3F_2\left[\begin{matrix}\alpha, & n_{\text{red}}, & n_{\text{ppl}} \\ & n_{\text{red}}+n_{\text{grey}}, & n_{\text{ppl}}+n_{\text{blue}}\end{matrix}; \beta(z-1)\right] \\
&= P\left(E_{m_2}\right)
\end{aligned}
$$

##### 3.5 The multi-state model for mRNAs by Zhou and Liu [13]

Let $s_0 = m_1 + i_1$, $s_1 = m_1 + i_1 + j_1$, $s_2 = m_1 + i_1 + j_1 + j_2$, ..., $s_k = m_1 + i_1 + \sum_{q=1}^{k} j_q$. Also, let $t_0 = i_1$, $t_1 = i_1 + j_1$, $t_2 = i_1 + j_1 + j_2$, ..., $t_k = i_1 + \sum_{q=1}^{k} j_q$ and $u_0 = 0$, $u_1 = j_1$, $u_2 = j_1 + j_2$, ..., $u_k = \sum_{q=1}^{k} j_q$. Finally, let $n_{1,q}$ and $n_{2,q}$ be the numbers of recipient urns of color 1 and color 2 in the $q^{\text{th}}$ layer.

$$
\begin{aligned}
P\left(E_{\text{urn},m_1}\right) &= P\left(\bullet_{m_1} \mapsto n_{\text{red}} | \mu, n_{\text{red}}, n_{\text{grey}}, n_{1,1}, n_{2,1}, n_{1,2}, n_{2,2}, n_{1,3}, n_{2,3}, ..., n_{1,\text{k}}, n_{2,\text{k}}, \right) \\
&= \sum_{i_1=0}^{\infty} \sum_{j_1=0}^{\infty} \sum_{j_2=0}^{\infty} ... \sum_{j_k=0}^{\infty} \text{Pois}\left(\bullet_{s_k} | \mu\right) \times \\
&\qquad \text{NH}\left(\bullet_{s_{k-1}} \mapsto n_{1,\text{k}} | \bullet_{s_k} \mapsto \{n_{1,\text{k}}, n_{2,\text{k}}\}\right) \times \\
&\qquad \text{NH}\left(\bullet_{s_{k-2}} \mapsto n_{1,\text{k-1}} | \bullet_{s_{k-1}} \mapsto \{n_{1,\text{k-1}}, n_{2,\text{k-1}}\}\right) \times \\
&\qquad \text{NH}\left(\bullet_{s_{k-3}} \mapsto n_{1,\text{k-2}} | \bullet_{s_{k-2}} \mapsto \{n_{1,\text{k-2}}, n_{2,\text{k-2}}\}\right) \times \\
&\qquad ... \times \\
&\qquad \text{NH}\left(\bullet_{s_0} \mapsto n_{1,1} | \bullet_{s_1} \mapsto \{n_{1,1}, n_{2,1}\}\right) \times \\
&\qquad \text{NH}\left(\bullet_{m_1} \mapsto n_{\text{red}} | \bullet_{m_1+i_1} \mapsto \{n_{\text{red}}, n_{\text{grey}}\}\right) \\
&= \sum_{i_1=0}^{\infty} \sum_{j_1=0}^{\infty} \sum_{j_2=0}^{\infty} ... \sum_{j_k=0}^{\infty} \frac{\binom{n_{\text{red}}+m_1-1}{m_1}\binom{n_{\text{grey}}+i_1-1}{i_1}}{\binom{n_{\text{red}}+n_{\text{grey}}+m_1+i_1-1}{m_1+i_1}} \prod_{q=1}^{k} \frac{\binom{n_{1,\text{q}}+s_{q-1}-1}{s_{q-1}}\binom{n_{2,\text{q}}+j_q-1}{j_q}}{\binom{n_{1,\text{q}}+n_{2,\text{q}}+s_q-1}{s_q}} \cdot \frac{e^{-\mu}\mu^{s_k}}{s_k!} \\
&= \frac{e^{-\mu}\mu^{m_1}}{m_1!} \sum_{i_1=0}^{\infty} \sum_{j_1=0}^{\infty} \sum_{j_2=0}^{\infty} ... \sum_{j_k=0}^{\infty} \frac{\left(n_{\text{red}}\right)_{m_1}\left(n_{\text{grey}}\right)_{i_1}}{\left(n_{\text{red}}+n_{\text{grey}}\right)_{m_1+i_1}} \frac{\mu^{i_1}}{i_1!} \prod_{q=1}^{k} \frac{\left(n_{1,\text{q}}\right)_{s_{q-1}}\left(n_{2,\text{q}}\right)_{j_q}}{\left(n_{1,\text{q}}+n_{2,\text{q}}\right)_{s_q}} \cdot \frac{\mu^{j_q}}{j_q!} \\
&\qquad\qquad\qquad\qquad\qquad\qquad\qquad\qquad \left[\text{Using } (a)_{x+y} = (a)_x\,(a+x)_y\right] \\
&= \frac{e^{-\mu}\mu^{m_1}}{m_1!} \frac{\left(n_{\text{red}}\right)_{m_1}}{\left(n_{\text{red}}+n_{\text{grey}}\right)_{m_1}} \left\{\prod_{q=1}^{k} \frac{\left(n_{1,\text{q}}\right)_{m_1}}{\left(n_{1,\text{q}}+n_{2,\text{q}}\right)_{m_1}}\right\} \cdot \\
&\qquad \sum_{i_1=0}^{\infty} \sum_{j_1=0}^{\infty} \sum_{j_2=0}^{\infty} ... \sum_{j_k=0}^{\infty} \frac{\left(n_{\text{grey}}\right)_{i_1}}{\left(n_{\text{red}}+n_{\text{grey}}+m_1\right)_{i_1}} \frac{\mu^{i_1}}{i_1!} \left\{\prod_{q=1}^{k} \frac{\left(n_{1,\text{q}}+m_1\right)_{t_{q-1}}\left(n_{2,\text{q}}\right)_{j_q}}{\left(n_{1,\text{q}}+n_{2,\text{q}}+m_1\right)_{t_q}} \cdot \frac{\mu^{j_q}}{j_q!}\right\} \\
&= \frac{\mu^{m_1}}{m_1!} \frac{\left(n_{\text{red}}\right)_{m_1}}{\left(n_{\text{red}}+n_{\text{grey}}\right)_{m_1}} \left\{\prod_{q=1}^{k} \frac{\left(n_{1,\text{q}}\right)_{m_1}}{\left(n_{1,\text{q}}+n_{2,\text{q}}\right)_{m_1}}\right\} \cdot \\
&\qquad e^{-\mu} \sum_{j_1=0}^{\infty} \sum_{j_2=0}^{\infty} ... \sum_{j_k=0}^{\infty} \left\{\prod_{q=1}^{k} \frac{\left(n_{2,\text{q}}\right)_{j_q}\left(n_{1,\text{q}}+m_1\right)_{u_{q-1}}}{\left(n_{1,\text{q}}+n_{2,\text{q}}+m_1\right)_{u_q}} \frac{\mu^{j_q}}{j_q!}\right\} \cdot \\
&\qquad\qquad \sum_{i_1=0}^{\infty} \left\{\prod_{q=1}^{k} \frac{\left(n_{1,\text{q}}+m_1+u_{q-1}\right)_{i_1}}{\left(n_{1,\text{q}}+n_{2,\text{q}}+m_1+u_q\right)_{i_1}}\right\} \frac{\left(n_{\text{grey}}\right)_{i_1}}{\left(n_{\text{red}}+n_{\text{grey}}+m_1\right)_{i_1}} \frac{\mu^{i_1}}{i_1!} \\
&= \frac{\mu^{m_1}}{m_1!} \frac{\left(n_{\text{red}}\right)_{m_1}}{\left(n_{\text{red}}+n_{\text{grey}}\right)_{m_1}} \left\{\prod_{q=1}^{k} \frac{\left(n_{1,\text{q}}\right)_{m_1}}{\left(n_{1,\text{q}}+n_{2,\text{q}}\right)_{m_1}}\right\} \cdot \\
&\qquad e^{-\mu} \sum_{j_1=0}^{\infty} \sum_{j_2=0}^{\infty} ... \sum_{j_k=0}^{\infty} \left\{\prod_{q=1}^{k} \frac{\left(n_{2,\text{q}}\right)_{j_q}\left(n_{1,\text{q}}+m_1\right)_{u_{q-1}}}{\left(n_{1,\text{q}}+n_{2,\text{q}}+m_1\right)_{u_q}} \frac{\mu^{j_q}}{j_q!}\right\} \cdot \\
&\qquad {}_{k+1}F_{k+1}\left[\begin{matrix} n_{1,1}+m_1+u_0, & n_{1,2}+m_1+u_1, & n_{1,3}+m_1+u_2, & ..., & n_{1,k}+m_1+u_{k-1}, & n_{\text{grey}} \\ n_{1,1}+n_{2,1}+m_1+u_1, & n_{1,2}+n_{2,2}+m_1+u_2, & n_{1,3}+n_{2,3}+m_1+u_3, & ..., & n_{1,k}+n_{2,k}+m_1+u_k, & n_{\text{red}}+n_{\text{grey}}+m_1 \end{matrix}; \mu\right] \\
&\qquad\qquad\qquad\qquad \left[\text{Using the Kummer-type transformation in Theorem 3 of Choudhary, 2020 [12]}\right] \\
&= \frac{\mu^{m_1}}{m_1!} \frac{\left(n_{\text{red}}\right)_{m_1}}{\left(n_{\text{red}}+n_{\text{grey}}\right)_{m_1}} \prod_{q=1}^{k} \frac{\left(n_{1,\text{q}}\right)_{m_1}}{\left(n_{1,\text{q}}+n_{2,\text{q}}\right)_{m_1}} \cdot \\
&\qquad {}_{k+1}F_{k+1}\left[\begin{matrix} n_{1,1}+m_1, & n_{1,2}+m_1, & n_{1,3}+m_1, & ..., & n_{1,k}+m_1, & n_{\text{red}}+m_1 \\ n_{1,1}+n_{2,1}+m_1, & n_{1,2}+n_{2,2}+m_1, & n_{1,3}+n_{2,3}+m_1, & ..., & n_{1,k}+n_{2,k}+m_1, & n_{\text{red}}+n_{\text{grey}}+m_1 \end{matrix}; -\mu\right] \\
&= \frac{1}{m_1!} \frac{d^{m_1}}{dz^{m_1}} {}_{k+1}F_{k+1}\left[\begin{matrix} n_{1,1}, & n_{1,2}, & n_{1,3}, & ..., & n_{1,k}, & n_{\text{red}} \\ n_{1,1}+n_{2,1}, & n_{1,2}+n_{2,2}, & n_{1,3}+n_{2,3}, & ..., & n_{1,k}+n_{2,k}, & n_{\text{red}}+n_{\text{grey}} \end{matrix}; \mu\,(z-1)\right]\Bigg|_{z=0} \\
&= \text{Coefficient of } z^{m_1} \text{ in the Maclaurin series expansion of} \\
&\qquad {}_{k+1}F_{k+1}\left[\begin{matrix} n_{1,1}, & n_{1,2}, & ..., & n_{1,k}, & n_{\text{red}} \\ n_{1,1}+n_{2,1}, & n_{1,2}+n_{2,2}, & ..., & n_{1,k}+n_{2,k}, & n_{\text{red}}+n_{\text{grey}} \end{matrix}; \mu\,(z-1)\right] \\
&= P\left(E_{m_1}\right)
\end{aligned}
$$

##### 3.6 The multi-state model for proteins

Let $s_0 = m_2 + i_2$, $s_1 = m_2 + i_2 + j_1$, $s_2 = m_2 + i_2 + j_1 + j_2$, ..., $s_k = m_2 + i_2 + \sum_{q=1}^{k} j_q$. Also, let $t_0 = i_2$, $t_1 = i_2 + j_1$, $t_2 = i_2 + j_1 + j_2$, ..., $t_k = i_2 + \sum_{q=1}^{k} j_q$ and $u_0 = 0$, $u_1 = j_1$, $u_2 = j_1 + j_2$, ..., $u_k = \sum_{q=1}^{k} j_q$. Finally, let $n_{1,q}$ and $n_{2,q}$ be the numbers of recipient urns of color 1 and color 2 in the $q^{\text{th}}$ layer.

$$
\begin{aligned}
P\left(E_{\text{urn},m_2}\right) &= P\left(\bullet_{m_2} \mapsto n_{\text{red}} | \alpha, \beta, n_{\text{red}}, n_{\text{grey}}, n_{1,1}, n_{2,1}, n_{1,2}, n_{2,2}, n_{1,3}, n_{2,3}, ..., n_{1,\text{k}}, n_{2,\text{k}}, \right) \\
&= \sum_{i_2=0}^{\infty} \sum_{j_1=0}^{\infty} \sum_{j_2=0}^{\infty} ... \sum_{j_k=0}^{\infty} \text{NB}\left(\bullet_{s_k} | \alpha, \beta\right) \times \\
&\qquad \text{NH}\left(\bullet_{s_{k-1}} \mapsto n_{1,\text{k}} | \bullet_{s_k} \mapsto \{n_{1,\text{k}}, n_{2,\text{k}}\}\right) \times \\
&\qquad \text{NH}\left(\bullet_{s_{k-2}} \mapsto n_{1,\text{k-1}} | \bullet_{s_{k-1}} \mapsto \{n_{1,\text{k-1}}, n_{2,\text{k-1}}\}\right) \times \\
&\qquad \text{NH}\left(\bullet_{s_{k-3}} \mapsto n_{1,\text{k-2}} | \bullet_{s_{k-2}} \mapsto \{n_{1,\text{k-2}}, n_{2,\text{k-2}}\}\right) \times \\
&\qquad ... \times \\
&\qquad \text{NH}\left(\bullet_{s_0} \mapsto n_{1,1} | \bullet_{s_1} \mapsto \{n_{1,1}, n_{2,1}\}\right) \times \\
&\qquad \text{NH}\left(\bullet_{m_2} \mapsto n_{\text{red}} | \bullet_{m_2+i_2} \mapsto \{n_{\text{red}}, n_{\text{grey}}\}\right) \\
&= \sum_{i_2=0}^{\infty} \sum_{j_1=0}^{\infty} \sum_{j_2=0}^{\infty} ... \sum_{j_k=0}^{\infty} \frac{\binom{n_{\text{red}}+m_2-1}{m_2}\binom{n_{\text{grey}}+i_2-1}{i_2}}{\binom{n_{\text{red}}+n_{\text{grey}}+m_2+i_2-1}{m_2+i_2}} \prod_{q=1}^{k} \frac{\binom{n_{1,\text{q}}+s_{q-1}-1}{s_{q-1}}\binom{n_{2,\text{q}}+j_q-1}{j_q}}{\binom{n_{1,\text{q}}+n_{2,\text{q}}+s_q-1}{s_q}} \cdot \binom{\alpha + s_k - 1}{s_k} \left(\frac{1}{1+\beta}\right)^{\alpha} \left(\frac{\beta}{1+\beta}\right)^{s_k} \\
&= \frac{\beta^{m_2}}{(1+\beta)^{\alpha+m_2} m_2!} \sum_{i_2=0}^{\infty} \sum_{j_1=0}^{\infty} \sum_{j_2=0}^{\infty} ... \sum_{j_k=0}^{\infty} \frac{(\alpha)_{s_k} (n_{\text{red}})_{m_2} (n_{\text{grey}})_{i_2}}{(n_{\text{red}} + n_{\text{grey}})_{m_2+i_2}} \frac{\beta^{i_2}}{(1+\beta)^{i_2} i_2!} \prod_{q=1}^{k} \frac{(n_{1,\text{q}})_{s_{q-1}} (n_{2,\text{q}})_{j_q}}{(n_{1,\text{q}} + n_{2,\text{q}})_{s_q}} \cdot \frac{1}{j_q!} \left(\frac{\beta}{1+\beta}\right)^{j_q} \\
&= \frac{\beta^{m_2}}{(1+\beta)^{\alpha+m_2} m_2!} \frac{(\alpha)_{m_2} (n_{\text{red}})_{m_2}}{(n_{\text{red}} + n_{\text{grey}})_{m_2}} \left\{\prod_{q=1}^{k} \frac{(n_{1,\text{q}})_{m_2}}{(n_{1,\text{q}} + n_{2,\text{q}})_{m_2}}\right\} \cdot \\
&\qquad \sum_{i_2=0}^{\infty} \sum_{j_1=0}^{\infty} \sum_{j_2=0}^{\infty} ... \sum_{j_k=0}^{\infty} \frac{(\alpha+m_2)_{t_k} (n_{\text{grey}})_{i_2}}{(n_{\text{red}} + n_{\text{grey}} + m_2)_{i_2}} \frac{\beta^{i_2}}{(1+\beta)^{i_2} i_2!} \left\{\prod_{q=1}^{k} \frac{(n_{1,\text{q}} + m_2)_{t_{q-1}} (n_{2,\text{q}})_{j_q}}{(n_{1,\text{q}} + n_{2,\text{q}} + m_2)_{t_q}} \cdot \frac{1}{j_q!} \left(\frac{\beta}{1+\beta}\right)^{j_q}\right\} \\
&= \frac{\beta^{m_2}}{m_2!} \frac{(\alpha)_{m_2} (n_{\text{red}})_{m_2}}{(n_{\text{red}} + n_{\text{grey}})_{m_2}} \left\{\prod_{q=1}^{k} \frac{(n_{1,\text{q}})_{m_2}}{(n_{1,\text{q}} + n_{2,\text{q}})_{m_2}}\right\} \cdot \\
&\qquad \left(\frac{1}{1+\beta}\right)^{\alpha+m_2} \sum_{j_1=0}^{\infty} \sum_{j_2=0}^{\infty} ... \sum_{j_k=0}^{\infty} (\alpha+m_2)_{u_k} \left\{\prod_{q=1}^{k} \frac{(n_{2,\text{q}})_{j_q} (n_{1,\text{q}} + m_2)_{u_{q-1}}}{(n_{1,\text{q}} + n_{2,\text{q}} + m_2)_{u_q}} \frac{1}{j_q!} \left(\frac{\beta}{1+\beta}\right)^{j_q}\right\} \cdot \\
&\qquad \sum_{i_2=0}^{\infty} (\alpha + m_2 + u_k)_{i_2} \left\{\prod_{q=1}^{k} \frac{(n_{1,\text{q}} + m_2 + u_{q-1})_{i_2}}{(n_{1,\text{q}} + n_{2,\text{q}} + m_2 + u_q)_{i_2}}\right\} \frac{(n_{\text{grey}})_{i_2}}{(n_{\text{red}} + n_{\text{grey}} + m_2)_{i_2}} \frac{\beta^{i_2}}{(1+\beta)^{i_2} i_2!} \\
&= \frac{\beta^{m_2}}{m_2!} \frac{(\alpha)_{m_2} (n_{\text{red}})_{m_2}}{(n_{\text{red}} + n_{\text{grey}})_{m_2}} \left\{\prod_{q=1}^{k} \frac{(n_{1,\text{q}})_{m_2}}{(n_{1,\text{q}} + n_{2,\text{q}})_{m_2}}\right\} \cdot \\
&\qquad \left(\frac{1}{1+\beta}\right)^{\alpha+m_2} \sum_{j_1=0}^{\infty} \sum_{j_2=0}^{\infty} ... \sum_{j_k=0}^{\infty} (\alpha+m_2)_{u_k} \left\{\prod_{q=1}^{k} \frac{(n_{2,\text{q}})_{j_q} (n_{1,\text{q}} + m_2)_{u_{q-1}}}{(n_{1,\text{q}} + n_{2,\text{q}} + m_2)_{u_q}} \frac{1}{j_q!} \left(\frac{\beta}{1+\beta}\right)^{j_q}\right\} \cdot \\
&\qquad {}_{k+2}F_{k+1}\left[\begin{matrix} \alpha+m_2+u_k, & n_{1,1}+m_2+u_0, & n_{1,2}+m_2+u_1, & ..., & n_{1,k}+m_2+u_{k-1}, & n_{\text{grey}} \\ & n_{1,1}+n_{2,1}+m_2+u_1, & n_{1,2}+n_{2,2}+m_2+u_2, & ..., & n_{1,k}+n_{2,k}+m_2+u_k, & n_{\text{red}}+n_{\text{grey}}+m_2 \end{matrix}; \frac{\beta}{1+\beta}\right]
\end{aligned}
$$

[Using the Euler-type transformation in Theorem 4 of Choudhary, 2020 [12]]

$$
\begin{aligned}
&= \frac{\beta^{m_2}}{m_2!} \frac{(\alpha)_{m_2} (n_{\text{red}})_{m_2}}{(n_{\text{red}} + n_{\text{grey}})_{m_2}} \prod_{q=1}^{k} \frac{(n_{1,\text{q}})_{m_2}}{(n_{1,\text{q}} + n_{2,\text{q}})_{m_2}} \cdot \\
&\qquad {}_{k+2}F_{k+1}\left[\begin{matrix} \alpha+m_2, & n_{1,1}+m_2, & n_{1,2}+m_2, & n_{1,3}+m_2, & ..., & n_{1,k}+m_2, & n_{\text{red}}+m_2 \\ & n_{1,1}+n_{2,1}+m_2, & n_{1,2}+n_{2,2}+m_2, & n_{1,3}+n_{2,3}+m_2, & ..., & n_{1,k}+n_{2,k}+m_2, & n_{\text{red}}+n_{\text{grey}}+m_2 \end{matrix}; -\beta\right] \\
&= \frac{1}{m_2!} \frac{d^{m_2}}{dz^{m_2}} {}_{k+2}F_{k+1}\left[\begin{matrix} \alpha, & n_{1,1}, & n_{1,2}, & n_{1,3}, & ..., & n_{1,k}, & n_{\text{red}} \\ & n_{1,1}+n_{2,1}, & n_{1,2}+n_{2,2}, & n_{1,3}+n_{2,3}, & ..., & n_{1,k}+n_{2,k}, & n_{\text{red}}+n_{\text{grey}} \end{matrix}; \beta(z-1)\right]\Bigg|_{z=0} \\
&= \text{Coefficient of } z^{m_2} \text{ in the Maclaurin series expansion of} \\
&\qquad {}_{k+2}F_{k+1}\left[\begin{matrix} \alpha, & n_{1,1}, & n_{1,2}, & ..., & n_{1,k}, & n_{\text{red}} \\ & n_{1,1}+n_{2,1}, & n_{1,2}+n_{2,2}, & ..., & n_{1,k}+n_{2,k}, & n_{\text{red}}+n_{\text{grey}} \end{matrix}; \beta(z-1)\right] \\
&= P\left(E_{m_2}\right)
\end{aligned}
$$

### 4 Deriving the probability of a given number of births using the inclusion-exclusion principle

In this section, we consider the Peccoud-Ycart model and provide an alternative way to derive the probability of $m_1$ transcriptions in an mRNA lifetime. Our urn scheme involves sampling $m_1 + i_1$ balls from the master urn with $i_1 > 0$, assigning them to $n_{\text{red}}$ red and $n_{\text{grey}}$ grey recipient urns, and writing the probability that *exactly* $m_1$ balls are assigned to the red recipient urns. One way to write this probability is directly using the negative hypergeometric distribution as described in Supplementary Section 2. Alternatively, we can obtain a series of probabilities for assigning *at least* $i_1$ balls to the grey urns, *at least* $i_1 + 1$ balls to the grey urns, ..., *at least* $i_1 + m_1$ balls to the grey urns and use these in conjunction with the inclusion-exclusion principle to derive the probability of *exactly* $m_1$ assignments to the red urns. While this is a less straightforward method, we use it to derive the probability distribution for mRNAs in the limit $n_{\text{red}} \gg n_{\text{grey}}$, which reveals that the transcriptional dynamics for highly expressed genes are more accurately described in terms of transcriptional lapses than in terms of transcriptional bursts.

Given that sampling from the master urn has yielded $m_1 + i_1$ black balls, we divide the sample into sets of $m_1$ and $i_1$ balls. This is doable in $\binom{m_1+i_1}{i_1}$ ways. Next, the probability that at least the $i_1$ balls are assigned to the grey urns is

$$\frac{\binom{n_{\text{grey}}+i_1-1}{i_1}}{\binom{n_{\text{red}}+n_{\text{grey}}+i_1-1}{i_1}}.$$

After assigning $i_1$ balls to the grey urns, we assign the remaining balls to the recipient urns in any possible way. Since the probability of all possible assignments of remaining balls is 1, it does not contribute any term to the above or subsequent expressions. Similarly, the probability that at least $1 + i_1$ balls are assigned to the grey urns is

$$\binom{m_1}{1}\frac{\binom{n_{\text{grey}}+1+i_1-1}{1+i_1}}{\binom{n_{\text{red}}+n_{\text{grey}}+1+i_1-1}{1+i_1}},$$

where $\binom{m_1}{1}$ is the number of ways to choose 1 ball out of the set of $m_1$ balls for assignment to the grey urns. In general, for $1 \leq k_1 \leq m_1$, the probability that at least $k_1 + i_1$ balls are assigned to the grey urns is

$$\binom{m_1}{k_1}\frac{\binom{n_{\text{grey}}+k_1+i_1-1}{k_i+i_1}}{\binom{n_{\text{red}}+n_{\text{grey}}+k_1+i_1-1}{k_1+i_1}}.$$

Next, to write the probability of more than $i_1$ assignments to the grey urns, we combine the probabilities for at least $k_1 + i_1$ assignments to grey urns for $1 \leq k_1 \leq m_1$ using the inclusion-exclusion principle to get

$$\sum_{k_1=1}^{m_1}(-1)^{k_1-1}\binom{m_1}{k_1}\frac{\binom{n_{\text{grey}}+k_1+i_1-1}{k_1+i_1}}{\binom{n_{\text{red}}+n_{\text{grey}}+k_1+i_1-1}{k_1+i_1}}. \tag{4}$$

Finally, to derive the probability of exactly $m_1$ assignments to the red urns given the sets of $m_1$ and $i_1$ balls, we deduct the above probability from the probability of assigning at least the $i_1$ balls to the grey urns to get

$$\frac{\binom{n_{\text{grey}}+i_1-1}{i_1}}{\binom{n_{\text{red}}+n_{\text{grey}}+i_1-1}{i_1}} - \sum_{k_1=1}^{m_1}(-1)^{k_1-1}\binom{m_1}{k_1}\frac{\binom{n_{\text{grey}}+k_1+i_1-1}{k_1+i_1}}{\binom{n_{\text{red}}+n_{\text{grey}}+k_1+i_1-1}{k_1+i_1}} = \sum_{k_1=0}^{m_1}(-1)^{k_1}\binom{m_1}{k_1}\frac{\binom{n_{\text{grey}}+k_1+i_1-1}{k_1+i_1}}{\binom{n_{\text{red}}+n_{\text{grey}}+k_1+i_1-1}{k_1+i_1}}. \tag{5}$$

We obtain the probability of interest by multiplying the result in Eq. 5 with the probability of sampling $m_1 + i_1$ balls from the master urn, the number of ways to divide them in sets of $m_1$ and $i_1$ balls and taking the sum over $i_1$ ranging from 0 to $\infty$, i.e., the probability of $m_1$ transcriptions is

$$P\left(E_{\text{urn},m_1}\right) = \sum_{i_1=0}^{\infty}\binom{m_1+i_1}{i_1}\frac{e^{-\mu}\mu^{m_1+i_1}}{(m_1+i_1)!}\sum_{k_1=0}^{m_1}(-1)^{k_1}\binom{m_1}{k_1}\frac{\binom{n_{\text{grey}}+k_1+i_1-1}{k_1+i_1}}{\binom{n_{\text{red}}+n_{\text{grey}}+k_1+i_1-1}{k_1+i_1}}. \tag{6}$$

It is helpful to check this result for 0, 1 and 2 transcriptions to ascertain its validity and grasp the intuition underlying the inclusion-exclusion principle. First, for $m_1 = 0$, Eq. 6 yields

$$\sum_{i_1=0}^{\infty}\frac{e^{-\mu}\mu^{i_1}}{(i_1)!}\frac{\binom{n_{\text{grey}}+i_1-1}{i_1}}{\binom{n_{\text{red}}+n_{\text{grey}}+i_1-1}{i_1}}, \tag{7}$$

which is the probability of any number of polymerase arrivals such that all of them are assigned to the grey urns. Further, it is the same as the probability for $m_1 = 0$ from the expression in the main text. Next, for $m_1 = 1$, Eq. 6 yields

$$\left\{\sum_{i_1=0}^{\infty}\binom{1+i_1}{i_1}\frac{e^{-\mu}\mu^{1+i_1}}{(1+i_1)!}\frac{\binom{n_{\text{grey}}+i_1-1}{i_1}}{\binom{n_{\text{red}}+n_{\text{grey}}+i_1-1}{i_1}}\right\} - \left\{\sum_{i_1=0}^{\infty}\binom{1+i_1}{i_1}\frac{e^{-\mu}\mu^{1+i_1}}{(1+i_1)!}\frac{\binom{n_{\text{grey}}+1+i_1-1}{1+i_1}}{\binom{n_{\text{red}}+n_{\text{grey}}+1+i_1-1}{1+i_1}}\right\}. \tag{8}$$

Here, the first term is the probability that one or more arrivals occur and at least all but one are assigned to the grey urns. By deducting the second term, which is the probability that all the arrivals are assigned to the grey urns yields the probability that exactly one arrival is assigned to the red urns. Finally, for $m_1 = 2$, Eq. 6 yields

$$\left\{\sum_{i_1=0}^{\infty}\binom{2+i_1}{i_1}\frac{e^{-\mu}\mu^{2+i_1}}{(2+i_1)!}\frac{\binom{n_{\text{grey}}+i_1-1}{i_1}}{\binom{n_{\text{red}}+n_{\text{grey}}+i_1-1}{i_1}}\right\} - \left\{2\sum_{i_1=0}^{\infty}\binom{2+i_1}{i_1}\frac{e^{-\mu}\mu^{2+i_1}}{(2+i_1)!}\frac{\binom{n_{\text{grey}}+1+i_1-1}{1+i_1}}{\binom{n_{\text{red}}+n_{\text{grey}}+1+i_1-1}{1+i_1}}\right\}$$
$$+\left\{\sum_{i_1=0}^{\infty}\binom{2+i_1}{i_1}\frac{e^{-\mu}\mu^{2+i_1}}{(2+i_1)!}\frac{\binom{n_{\text{grey}}+2+i_1-1}{2+i_1}}{\binom{n_{\text{red}}+n_{\text{grey}}+2+i_1-1}{2+i_1}}\right\}. \tag{9}$$

Here again, the first is the probability that two or more arrivals occur and at least all but two are assigned to the grey urns. Let us label the two arrivals in addition to $i_1$ as $j_1$ and $j_2$. Then, there are two ways to ensure that at least all but one arrivals are assigned to the grey urns — by mixing $j_1$ with $i_1$ arrivals and assigning them to the grey urns or by mixing $j_2$ with $i_1$ arrivals and assigning them to the grey urns. This contributes the factor of 2 outside the summation in the second pair of curly brackets. However, note that the probability of each of these two ways includes the probability of assigning both $j_1$ and $j_2$ to the grey urns. Hence, by deducting the probability of both from the first term, we effectively deduct the probability of assigning both $j_1$ and $j_2$ to the grey urns twice. Hence, we add it once to the expression as the third term, which represents the probability of all arrivals assigned to the grey urns. Finally, the complete expression represents the probability of exactly two arrivals assigned to the red urns. The same principle applies to $m_1 > 2$.

Eq. 6 can be written after rearranging terms as

$$\begin{aligned} P\left(E_{\mathrm{urn},m_1}\right) &= \frac{1}{m_1!}\sum_{k_1=0}^{m_1}\binom{m_1}{k_1}e^{-\mu}\mu^{m_1-k_1}\left(-\mu\right)^{k_1}\sum_{i_1=0}^{\infty}\frac{\left(n_{\mathrm{grey}}\right)_{k_1+i_1}}{\left(n_{\mathrm{red}}+n_{\mathrm{grey}}\right)_{k_1+i_1}}\frac{\mu^{i_1}}{i_1!} \\ &= \frac{1}{m_1!}\sum_{k_1=0}^{m_1}\binom{m_1}{k_1}\left\{e^{-\mu}\mu^{m_1-k_1}\right\}\left\{\frac{\left(n_{\mathrm{grey}}\right)_{k_1}}{\left(n_{\mathrm{red}}+n_{\mathrm{grey}}\right)_{k_1}}\left(-\mu\right)^{k_1}\sum_{i_1=0}^{\infty}\frac{\left(n_{\mathrm{grey}}+k_1\right)_{i_1}}{\left(n_{\mathrm{red}}+n_{\mathrm{grey}}+k_1\right)_{i_1}}\frac{\mu^{i_1}}{i_1!}\right\} \\ &= \frac{1}{m_1!}\sum_{k_1=0}^{m_1}\binom{m_1}{k_1}\left\{e^{-\mu}\mu^{m_1-k_1}\right\}\left\{\frac{\left(n_{\mathrm{grey}}\right)_{k_1}}{\left(n_{\mathrm{red}}+n_{\mathrm{grey}}\right)_{k_1}}\left(-\mu\right)^{k_1}{}_1F_1\left(\begin{matrix}n_{\mathrm{grey}}+k_1\\ n_{\mathrm{red}}+n_{\mathrm{grey}}+k_1\end{matrix};\mu\right)\right\}. \end{aligned} \tag{10}$$

In the above expression, the term in the first pair of curly brackets is the same as

$$\left.\frac{d^{m_1-k_1}}{dz^{m_1-k_1}}e^{\mu(z-1)}\right|_{z=0}$$

and the term in the second pair of curly brackets is the same as

$$\left.\frac{d^{k_1}}{dz^{k_1}}{}_1F_1\left[\begin{matrix}n_{\mathrm{grey}}\\ n_{\mathrm{red}}+n_{\mathrm{grey}}\end{matrix};-\mu\left(z-1\right)\right]\right|_{z=0}.$$

Indeed, Eq. 10 is the coefficient of $z^{m_1}$ in the Maclaurin series expansion of $e^{\mu(z-1)}{}_1F_1\left[\begin{smallmatrix}n_{\mathrm{grey}}\\ n_{\mathrm{red}}+n_{\mathrm{grey}}\end{smallmatrix};-\mu\left(z-1\right)\right]$, which can be obtained by applying the general Leibniz's rule for differentiation and evaluating the derivative at $z=0$. Hence, the generating function for this probability is given by $e^{\mu(z-1)}{}_1F_1\left[\begin{smallmatrix}n_{\mathrm{grey}}\\ n_{\mathrm{red}}+n_{\mathrm{grey}}\end{smallmatrix};-\mu\left(z-1\right)\right]$. A straightforward application of the Kummer transformation formula for confluent hypergeometric function of the first kind shows that this generating function is the same as ${}_1F_1\left[\begin{smallmatrix}n_{\mathrm{red}}\\ n_{\mathrm{red}}+n_{\mathrm{grey}}\end{smallmatrix};\mu\left(z-1\right)\right]$, which is the Peccoud-Ycart solution.

#### 5 Limiting forms of the exact solution of the Peccoud-Ycart model

In this section, we derive the limiting forms of the exact solution of the Peccoud-Ycart model (Fig. 2a) for activated and repressed systems. To this end, we work with the leaky two-state model (Fig. 2c) and write the solutions for the Peccoud-Ycart model as its special case. In the leaky two-state model, a gene switches from the leaky to the active state with a propensity $k_0$ and from the active to the leaky state with a propensity $k_1$. The propensity of transcription in the leaky state is given by a leakage factor, $\lambda$ times the propensity of transcription in the active state, $v_0$. The mRNA degrades with a propensity $d_0$. When $\lambda = 0$, the leaky two-state model reduces to the Peccoud-Ycart model.

##### 5.1 Master equations

First, we write the chemical master equations. At any time, the system is described by two variables, the mRNA copy number, $m$ and the gene state. We use $p_{m,0}$ to denote the probability of there being $m$ mRNAs when the gene is in the leaky state and $p_{m,1}$ to denote the probability of there being $m$ mRNAs when the gene is in the active state. Now, the master equations can be written as

$$\frac{dp_{m,0}}{dt} = (k_1 p_{m,1} - k_0 p_{m,0}) + d_0 \left[(m+1) p_{m+1,0} - m p_{m,0}\right] + v_0 \lambda \left(p_{m-1,0} - p_{m,0}\right), \tag{11}$$

$$\frac{dp_{m,1}}{dt} = (k_0 p_{m,0} - k_1 p_{m,1}) + d_0 \left[(m+1) p_{m+1,1} - m p_{m,1}\right] + v_0 \left(p_{m-1,1} - p_{m,1}\right). \tag{12}$$

We are interested in solving for the probability, $\tilde{p}_m$ that there will be $m$ mRNAs in a cell at stationary state. To this end, we put the derivatives with respect to time in Eqs. 11-12 as 0, denote the stationary state $p_{m,0}$ and $p_{m,1}$ as $\tilde{p}_{m,0}$ and $\tilde{p}_{m,1}$, respectively, and define $\tilde{p}_m = \tilde{p}_{m,0} + \tilde{p}_{m,1}$. Adding Eqs. 11-12 gives an equation for $\tilde{p}_m$ at stationary state —

$$0 = d_0 \left[(m+1) \tilde{p}_{m+1} - m \tilde{p}_m\right] + v_0 \lambda \left(\tilde{p}_{m-1} - \tilde{p}_m\right) + v_0 (1-\lambda) \left(\tilde{p}_{m-1,1} - \tilde{p}_{m,1}\right). \tag{13}$$

Let us define generating functions, $\tilde{g}_1(z) = \sum_m z^m \tilde{p}_{m,1}$ and $\tilde{g}(z) = \sum_m z^m \tilde{p}_m$. Note that being a generating function for a probability distribution, $\tilde{g}(1) = 1$. Additionally, let us define $n_{\text{red}} = k_0/d_0$, $n_{\text{grey}} = k_1/d_0$ and $\mu = v_0/d_0$. Now, in terms of $\tilde{g}_1$ and $\tilde{g}$, Eqs. 12-13 can be expressed at stationary state as

$$(z-1)\frac{d\tilde{g}_1}{dz} = n_{\text{red}}\tilde{g} - (n_{\text{red}} + n_{\text{grey}})\tilde{g}_1 + \mu (z-1)\tilde{g}_1, \tag{14}$$

$$(z-1)\frac{d\tilde{g}}{dz} = \mu\lambda (z-1)\tilde{g} + \mu (1-\lambda)(z-1)\tilde{g}_1. \tag{15}$$

Eliminating $\tilde{g}_1$ from Eqs. 14-15, we get a second order differential equation for $\tilde{g}$,

$$(z-1)\frac{d^2\tilde{g}}{dz^2} + \left[n_{\text{red}} + n_{\text{grey}} - \mu (z-1)(1+\lambda)\right]\frac{d\tilde{g}}{dz} - \mu\left[n_{\text{red}} + n_{\text{grey}}\lambda - \mu (z-1)\lambda\right]\tilde{g} = 0. \tag{16}$$

Using a linear transformation, $u = \mu (1-\lambda)(z-1)$ and defining $f(u) = \exp(-\lambda u/1-\lambda)\tilde{g}$, Eq. 16 becomes

$$u\frac{d^2 f}{du^2} + (n_{\text{red}} + n_{\text{grey}} - u)\frac{df}{du} - n_{\text{red}} f = 0. \tag{17}$$

Note that the boundary condition on generating function, $\tilde{g}(1) = 1$ implies $f(0) = 1$.

##### 5.2 Limiting distribution in the repressed case

For a repressed gene, $n_{\text{red}} \ll n_{\text{grey}}$. We define a perturbation parameter, $\epsilon = n_{\text{red}}/n_{\text{red}}+n_{\text{grey}}$. Upon multiplying both sides by $\epsilon/n_{\text{red}}$, Eq. 17 yields

$$\frac{\epsilon u}{n_{\text{red}}}\frac{d^2 f}{du^2} + \left(1 - \frac{\epsilon u}{n_{\text{red}}}\right)\frac{df}{du} - \epsilon f = 0. \tag{18}$$

In terms of the perturbation parameter, $f(u)$ can be described as a perturbation series —

$$f(u,\epsilon) = f_0(u) + \epsilon f_1(u) + \epsilon^2 f_2(u) + \mathcal{O}\left(\epsilon^3\right). \tag{19}$$

Note that the boundary condition on $f(0) = 1$ still holds, i.e., $f(0,\epsilon) = 1$. Using the perturbation series, Eq. 18 can be rewritten as

$$\frac{\epsilon u}{n_{\text{red}}}\left(\frac{d^2 f_0}{du^2} + \epsilon\frac{d^2 f_1}{du^2} + \mathcal{O}\left(\epsilon^2\right)\right) + \left(1 - \frac{\epsilon u}{n_{\text{red}}}\right)\left(\frac{df_0}{du} + \epsilon\frac{df_1}{du} + \epsilon^2\frac{df_2}{du} + \mathcal{O}\left(\epsilon^3\right)\right) - \epsilon\left(f_0 + \epsilon f_1 + \mathcal{O}\left(\epsilon^2\right)\right) = 0. \tag{20}$$

Collecting together terms of same order in $\epsilon$, we get

$$\frac{df_0}{du} = 0, \tag{21}$$

$$\frac{df_1}{du} = f_0 + \frac{u}{n_{\text{red}}}\frac{df_0}{du} - \frac{u}{n_{\text{red}}}\frac{d^2 f_0}{du^2}, \tag{22}$$

$$\frac{df_2}{du} = f_1 + \frac{u}{n_{\text{red}}}\frac{df_1}{du} - \frac{u}{n_{\text{red}}}\frac{d^2 f_1}{du^2}. \tag{23}$$

Eqs. 21-23 can be solved with the boundary condition, $f(0,\epsilon)=1$ which implies that $f_0(0)=1$ and $f_1(0)=f_2(0)=...=0$. On solving Eqs. 21-23 with these boundary conditions, we find

$$\frac{df_0}{du}=0 \implies f_0(u)=1, \tag{24}$$

$$\frac{df_1}{du}=1 \implies f_1(u)=u, \tag{25}$$

$$\frac{df_2}{du}=\left(\frac{n_{\text{red}}+1}{n_{\text{red}}}\right)u \implies f_2(u)=\left(\frac{n_{\text{red}}+1}{n_{\text{red}}}\right)\frac{u^2}{2}. \tag{26}$$

In general, the overall solution can be written as

$$f(u,\epsilon) = 1+\epsilon u+\epsilon^2\left(\frac{n_{\text{red}}+1}{n_{\text{red}}}\right)\frac{u^2}{2}+... \tag{27}$$

$$= 1+\frac{n_{\text{red}}}{n_{\text{red}}+n_{\text{grey}}}u+\frac{n_{\text{red}}(n_{\text{red}}+1)}{(n_{\text{red}}+n_{\text{grey}})^2}\frac{u^2}{2!}+... \tag{28}$$

$$\implies f(u) = \left(1-\frac{u}{n_{\text{red}}+n_{\text{grey}}}\right)^{-n_{\text{red}}}. \tag{29}$$

provided $\left|\frac{u}{n_{\text{red}}+n_{\text{grey}}}\right|<1$. Hence, the generating function for the probability distribution in the repressed condition can be given as

$$\tilde{g}(z) = e^{\mu\lambda(z-1)}\cdot[1-\beta_r(z-1)]^{-\alpha_r}, \tag{30}$$

where $\beta_r = {}^{\mu(1-\lambda)}/_{n_{\text{red}}+n_{\text{grey}}} \approx {}^{\mu(1-\lambda)}/_{n_{\text{grey}}}$ is the burst size and $\alpha_r = n_{\text{red}}$ is the burst frequency. In other words, mRNAs follow a Delaporte distribution, which is a convolution of Poisson and negative binomial distributions. A constant minimal rate of transcription, $v_0\lambda$ gives rise to the Poisson component while gene activation leads to bursty phases of transcription contributing the negative binomial part. If $\lambda=0$, i.e., for the Peccoud-Ycart model, this distribution reduces to a negative binomial distribution, $[1-{}^{\mu(z-1)}/_{n_{\text{grey}}}]^{-n_{\text{red}}}$.

##### 5.3 Limiting distribution in the activated case

For an activated gene, $n_{\text{grey}} \ll n_{\text{red}}$. We define a perturbation parameter, $\epsilon = {}^{n_{\text{grey}}}/_{n_{\text{red}}+n_{\text{grey}}}$. Using this parameter, Eq. 17 yields

$$\frac{\epsilon u}{n_{\text{grey}}}\frac{d^2f}{du^2}+\left(1-\frac{\epsilon u}{n_{\text{grey}}}\right)\frac{df}{du}-(1-\epsilon)f = 0. \tag{31}$$

In terms of the perturbation parameter, $f(u)$ can again be described as a perturbation series —

$$f(u,\epsilon) = f_0(u)+\epsilon f_1(u)+\epsilon^2 f_2(u)+\mathcal{O}(\epsilon^3). \tag{32}$$

The boundary condition on $f(0)=1$ implies $f(0,\epsilon)=1$. Using the perturbation series, Eq. 31 can be rewritten as

$$\frac{\epsilon u}{n_{\text{grey}}}\left(\frac{d^2f_0}{du^2}+\epsilon\frac{d^2f_1}{du^2}+\mathcal{O}(\epsilon^2)\right)+\left(1-\frac{\epsilon u}{n_{\text{grey}}}\right)\left(\frac{df_0}{du}+\epsilon\frac{df_1}{du}+\epsilon^2\frac{df_2}{du}+\mathcal{O}(\epsilon^3)\right)$$
$$-(1-\epsilon)\left(f_0+\epsilon f_1+\mathcal{O}(\epsilon^2)\right) = 0. \tag{33}$$

Collecting together terms of same order in $\epsilon$, we get

$$\frac{df_0}{du} = f_0, \tag{34}$$

$$\frac{df_1}{du}-f_1 = -f_0+\frac{u}{n_{\text{grey}}}\frac{df_0}{du}-\frac{u}{n_{\text{grey}}}\frac{d^2f_0}{du^2}, \tag{35}$$

$$\frac{df_2}{du}-f_2 = -f_1+\frac{u}{n_{\text{grey}}}\frac{df_1}{du}-\frac{u}{n_{\text{grey}}}\frac{d^2f_1}{du^2}. \tag{36}$$

Eqs. 34-36 can be solved with the boundary condition, $f(0,\epsilon)=1$ which implies that $f_0(0)=1$ and $f_1(0)=f_2(0)=...=0$. On solving Eqs. 34-36 with these boundary conditions, we find

$$\frac{df_0}{du}=f_0 \implies f_0(u)=e^u, \tag{37}$$

$$\frac{df_1}{du}-f_1=-e^u \implies f_1(u)=-ue^u, \tag{38}$$

$$\frac{df_2}{du}-f_2=\left(\frac{n_{\text{grey}}+1}{n_{\text{grey}}}\right)ue^u \implies f_2(u)=\left(\frac{n_{\text{grey}}+1}{n_{\text{grey}}}\right)\frac{u^2}{2}e^u. \tag{39}$$

In general, the overall solution can be written as

$$f(u,\epsilon) = e^{u} + \epsilon\left(-ue^{u}\right) + \epsilon^{2}\left(\frac{n_{\text{grey}}+1}{n_{\text{grey}}}\right)\frac{u^{2}}{2}e^{u} + ... \tag{40}$$

$$= e^{u}\left[1 - \frac{n_{\text{grey}}}{n_{\text{red}}+n_{\text{grey}}}u + \frac{n_{\text{grey}}\left(n_{\text{grey}}+1\right)}{\left(n_{\text{red}}+n_{\text{grey}}\right)^{2}}\frac{u^{2}}{2!} + ...\right] \tag{41}$$

$$\implies f(u) = e^{u}\left(1+\frac{u}{n_{\text{red}}+n_{\text{grey}}}\right)^{-n_{\text{grey}}} \tag{42}$$

provided $\left|\frac{u}{n_{\text{red}}+n_{\text{grey}}}\right| < 1$. Hence, the generating function for the probability distribution in the activated condition can be given as

$$\tilde{g}(z) = e^{\mu(z-1)} \cdot \left[1-\beta_a(z-1)\right]^{-\alpha_a}, \tag{43}$$

where $\beta_a = -{}^{\mu(1-\lambda)}/_{n_{\text{red}}+n_{\text{grey}}} \approx -{}^{\mu(1-\lambda)}/_{n_{\text{red}}}$ and $\alpha_a = n_{\text{grey}}$. Note that Eq. 43 is formally identical to Eq. 30 but $\beta_a$ is negative while $\beta_r$ is positive. A negative $\beta_a$ also implies that the mRNA distribution under activated condition is not strictly Delaporte distribution but similar to Delaporte distribution. When $\lambda = 0$, i.e., for the Peccoud-Ycart model, $\beta_a = -{}^{\mu}/_{n_{\text{red}}}$. The negative value of $\beta_a$ can be interpreted as the mean number of mRNAs that were not produced due to an occurrence of a transcriptional *lapse*, i.e., a short-lived phase of inactivation of the gene.

#### References

[1] O G Berg. “A model for the statistical fluctuations of protein numbers in a microbial population.” In: *Journal of Theoretical Biology* 71.4 (Apr. 1978), pp. 587–603. ISSN: 0022-5193.

[2] Norman L Johnson and Samuel Kotz. *Urn Models and Their Application.* John Wiley & Sons, 1977.

[3] Vahid Shahrezaei and Peter S Swain. “Analytical distributions for stochastic gene expression.” In: *Proceedings of the National Academy of Sciences* 105.45 (Nov. 2008), pp. 17256–61. ISSN: 1091-6490.

[4] Norman L Johnson, Adrienne W Kemp, and Samuel Kotz. *Univariate discrete distributions.* Vol. 444. John Wiley & Sons, 2005.

[5] “Negative hypergeometric distribution”. In: Encyclopedia of Mathematics. URL: `http://www.encyclopediaofmath.org/index.php?title=Negative_hypergeometric_distribution&oldid=41612`.

[6] William C Guenther. “The inverse hypergeometric —a useful model”. In: *Statistica Neerlandica* 29.4 (1975), pp. 129–144.

[7] Jean Peccoud and Bernard Ycart. “Markovian modeling of gene product synthesis”. In: *Theoretical Population Biology* 48 (1995), pp. 222–234.

[8] Milton Abramowitz and Irene A Stegun. *Handbook of mathematical functions with formulas, graphs, and mathematical tables.* 1970.

[9] Zhixing Cao et al. “Multi-scale bursting in stochastic gene expression”. In: *bioRxiv* (2019), p. 717199.

[10] Rajesh Karmakar. “Conversion of graded to binary response in an activator-repressor system”. In: *Physical Review E* 81.2 (2010), p. 021905.

[11] RB Paris. “A Kummer-type transformation for a ${}_2F_2$ hypergeometric function”. In: *Journal of Computational and Applied Mathematics* 173.2 (2005), pp. 379–382.

[12] Krishna Choudhary. “Addition formulas for the ${}_pF_p$ and ${}_{p+1}F_p$ generalized hypergeometric functions with arbitrary parameters and their Kummer- and Euler-type transformations”. In: *arXiv preprint arXiv:2001.03815* (2020).

[13] Tianshou Zhou and Tuoqi Liu. “Quantitative analysis of gene expression systems”. In: *Quantitative Biology* 3.4 (2015), pp. 168–181.
